# Independent major-effect loci and limited trait associations characterise warning colour variation across sexes and life stages in the wood tiger moth (*Arctia plantaginis*)

**DOI:** 10.64898/2026.06.03.729769

**Authors:** Eva L. Koch, Melanie N. Brien, Yingguang Frank Chan, Marek Kučka, Cristina Ottocento, Chiara De Pasqual, Eetu Selenius, Ossi Nokelainen, Juan A. Galarza, Sandra Winters, Johanna Mappes, Chris D. Jiggins

## Abstract

Variation in warning colouration is common, despite strong positive frequency-dependent selection favouring the most common morph, and may arise from spatially or temporally variable selection or associations with other traits. We investigated the maintenance of warning colour variation in the wood tiger moth, *Arctia plantaginis,* which shows sexual dimorphism in colouration: two distinct male morphs controlled by a single locus, and continuous variation in female hindwing colour and larval warning signal size. Using whole-genome sequencing of 657 individuals from a laboratory stock derived from a wild polymorphic population, we examined the genetic basis of multiple life-history traits, chemical defence, pheromones, and male activity. We specifically tested for associations with male colour and aimed to identify the genetic basis of female and larval colour variation. We detected separate major-effect loci for female colour and larval signal size, which were different from the male colour locus. Hindwing melanism was the only trait associated with male colour, mapping to a nearby major-effect locus and showing a link to reduced activity at higher temperatures. Female colour was phenotypically correlated with pheromone amount. However, apart from melanism, none of the other traits measured showed an association with any of the loci controlling warning colours and we found no evidence that male colour is part of a ‘supergene’ or complex co-adapted phenotype. Instead, our results indicate largely independent genetic architectures for warning colours suggesting that variable selection rather than genetic associations may contribute to the maintenance of colour polymorphisms in this species.

## Introduction

Variation in warning colouration is widespread among aposematic species (Briolat et al. 2019), yet its persistence remains puzzling. Predators are expected to learn and associate the most common warning signal with unpalatability more quickly, leading to positive frequency-dependent selection and evolution of a consistent warning signal. It was indeed shown that the selection pressures for consistent warning can be strong (Mallet and Barton 1989), sometimes driving convergence in colour patterns even across species (Joron et al. 1999) However, predator-driven selection can vary spatially (Joron et al. 1999; Chouteau et al. 2016) and seasonally (Mappes et al. 2014). The selective advantage of a particular colour pattern may change depending on predation risk, the composition of the local predator community, the presence of other aposematic species, habitat complexity, and variation in sexual selection (reviewed in McLean and Stuart-Fox (2014)). Moreover, colour is often subject to multiple selection pressures. It can function as a cue in mate attraction and be shaped by sexual selection (Wellenreuther et al. 2014), play a role in thermoregulation and immune responses (True 2003), and incur costs associated with the production of colour pigments (Graham et al. 1980). Trade-offs between these costs and benefits may vary across environments and over time, creating a complex mosaic of selection pressures that help maintain variation in warning colouration.

Another interesting aspect of colour polymorphisms is their frequent correlation with other traits (McKinnon and Pierotti 2010). Morphs may differ not only in colour but also in other characteristics linked to alternative strategies for maximizing fitness (Oliveira et al. 2008). As a result, selection favours different adaptive trait combinations in different morphs. For example, in ruffs, three male morphs differ in plumage, behaviour, body size, and testis volume (Küpper et al. 2016). In white-throated sparrows, colour morphs vary in parental care, and extra-pair mating attempts (Tuttle et al. 2016), while in side-blotched lizards, male colour is associated with distinct reproductive strategies: mate guarding, sneaking, and aggression (Zamudio and Sinervo 2000). Different alternative strategies can be maintained through negative frequency-dependent selection (Zamudio and Sinervo 2000), disassortative mating (Tuttle et al. 2016), or temporal fluctuations in morph-specific success (Olsson et al. 2007). Warning colour could thus be part of a complex phenotype in species where polymorphisms persist. The genetic basis of complex phenotypes are often supergenes, clusters of tightly linked genes with little or no recombination (Schwander et al. 2014). Among the best-studied cases are chromosomal inversions, which prevent recombination between alternative arrangements in heterozygous individuals and can thus maintain sets of adaptive alleles. In some cases, these inversions can drive alternative reproductive strategies and control colour (Lamichhaney et al. 2016; Tuttle et al. 2016). Thus, in addition to spatial and temporal variation in selection, associations between colour and other traits may help explain the maintenance of polymorphisms.

In this study, we investigate colour polymorphisms in the wood tiger moth *Arctia plantaginis* (former *Parasemia plantaginis*, Arctiinae (Rönkä et al. 2016)). *A. plantaginis* is an aposematic species. Adults release defensive fluids when attacked (Rojas et al. 2017; Burdfield-Steel et al. 2018; Rojas et al. 2019) and use hindwing colour to signal their unpalatability to avian predators (Lindstedt et al. 2011; Nokelainen et al. 2012). Males exhibit either distinct white or yellow hindwings and polymorphisms persist within Finnish populations (Galarza et al. 2014). Inheritance patterns indicated colour to be controlled by a single locus with two alleles, with the white allele being dominant (Nokelainen et al. 2012; Nokelainen et al. 2022a). Contrary to the expectation that male colour might be part of a complex phenotype controlled by a supergene, the genetic basis has been identified as a narrow (100 kb) genomic region, resulting from a gene duplication event. Male colour is controlled by a copy of the *yellow-e* gene within this region, named *valkea* (Brien et al. 2023). However, supergenes may not always be necessary to maintain complex alternative phenotypes (Flanagan and Alonzo 2024), and complex phenotypes can also be controlled by a single genomic region (Thompson et al. 2020).

In *A. plantaginis*, pleiotropic effects of the duplicated region on other traits may still contribute to the maintenance of the polymorphism. Loci with pleiotropic effects have been detected in other species. In Gouldian finches, a sex-linked colour polymorphism is associated with differences in social dominance, sperm morphometry, and sex allocation, and controlled by a small (72 kb) genomic region (Kim et al. 2019). In Lepidoptera, the gene *aristaless* can influence both wing colour and mate choice (Westerman et al. 2018). Associations between colour and other traits may also arise from the costs of pigment synthesis. In *Colias* butterflies, wing colour is controlled by a single locus and correlated with growth and reproduction due to a resource allocation trade-off: white females develop faster and produce more eggs than those with orange hindwings. The persistence of two colour morphs is maintained by multiple factors, including temperature, increased viral sensitivity in white morphs, and mate preferences (Woronik et al. 2019).

Many studies on *A. plantaginis* have focused on the discrete male hindwing colour morphs. However, aposematism in this species is multifaceted, occurring in both sexes and larvae. Unlike males, females exhibit continuous variation in hindwing colour (Hegna et al. 2015), ranging from yellow to red, with red providing the strongest deterrent effect against predators (Lindstedt et al. 2011; Rönkä et al. 2018). Larvae are also unprofitable (Lindstedt et al. 2008), and both sexes display an orange patch on their black bodies, which varies in size and contributes to predator deterrence (Lindstedt et al. 2008; Nielsen and Mappes 2020). These traits show some degree of plasticity (Ojala et al. 2007; Lindstedt et al. 2010; Galarza et al. 2019; Almeida et al. 2021), but also substantial genetic variation (Ojala et al. 2007; Lindstedt et al. 2009; Lindstedt et al. 2016; Koch et al. 2024) but their genetic basis remains unknown. Studies suggest that they are largely independent, with no genetic correlation and little to no phenotypic correlation between larval, female and male warning colouration (Lindstedt et al. 2010; Lindstedt et al. 2020; Nokelainen et al. 2022a; Koch et al. 2024).

In this study, we investigate the maintenance of variation in warning colours in *A. plantaginis,* specifically exploring whether male colour is part of a complex phenotype. We further aim to identify the genetic basis of female and larval colour variation and examine how diverse traits under selection including warning colours are correlated and linked due to shared resources or a common genetic basis to better understand potential trade-offs and constraints in this species. We address the following key questions: Is male hindwing colour part of a complex phenotype, i.e. a suite of multiple traits that are correlated with each other and may contribute to different adaptive strategies within a population? What is the genetic architecture of other colour traits in this species? Does colour variation correlate with other fitness-relevant traits such as chemical defence or pheromones? How are these different traits related in general?

To answer these questions, we first tested for associations between male colour and other traits in a laboratory-raised population maintained for over 20 generations. Using whole-genome sequencing of a subset of these individuals, we then investigated the genetic basis of a broad range of traits, including size, developmental time, activity, and the composition of defensive fluids and pheromones, to determine whether these traits are influenced by the male colour locus. Additionally, these data also allow us to test for phenotypic associations between diverse traits under selection, highlighting potential costs and trade-offs. By addressing these questions, our study provides insights into how genetic architecture and different selection pressures may have shaped warning colour variation in *A. plantaginis*.

## Methods

### Animal rearing

Animals came from a laboratory population of Finnish ancestry that had been maintained for 25 generations. Details on the maintenance of this population can be found in De Pasqual et al. (2022). Briefly, the initial generations were established in 2013 using adult wild Finnish individuals. In nature, only one generation occurs per year, with all larvae overwintering and completing their development the following year. Under laboratory conditions, three generations can be raised per year, with two completing development without diapause. To set up the next generation, one adult male and one adult female were placed in containers for mating and egg laying. Each individual was only mated once. Hatched larvae stayed in these containers for the first weeks of their development before being separated and kept in groups of approximately 30 individuals. Developing larvae were fed daily with wild dandelion and checked for pupation. Pupae were then weighed and kept individually until adulthood. Dates of hatching, pupation, and eclosion were recorded for all individuals, as well as pupa weight, male hindwing colour, and their melanisation type (plus or minus, see Figure 1F). Larval time refers to the number of days from hatching to pupation. The colour of adult females was measured by visually matching their hindwings against a colour scale ranging from one (yellow) to six (red).

**Figure 1.**
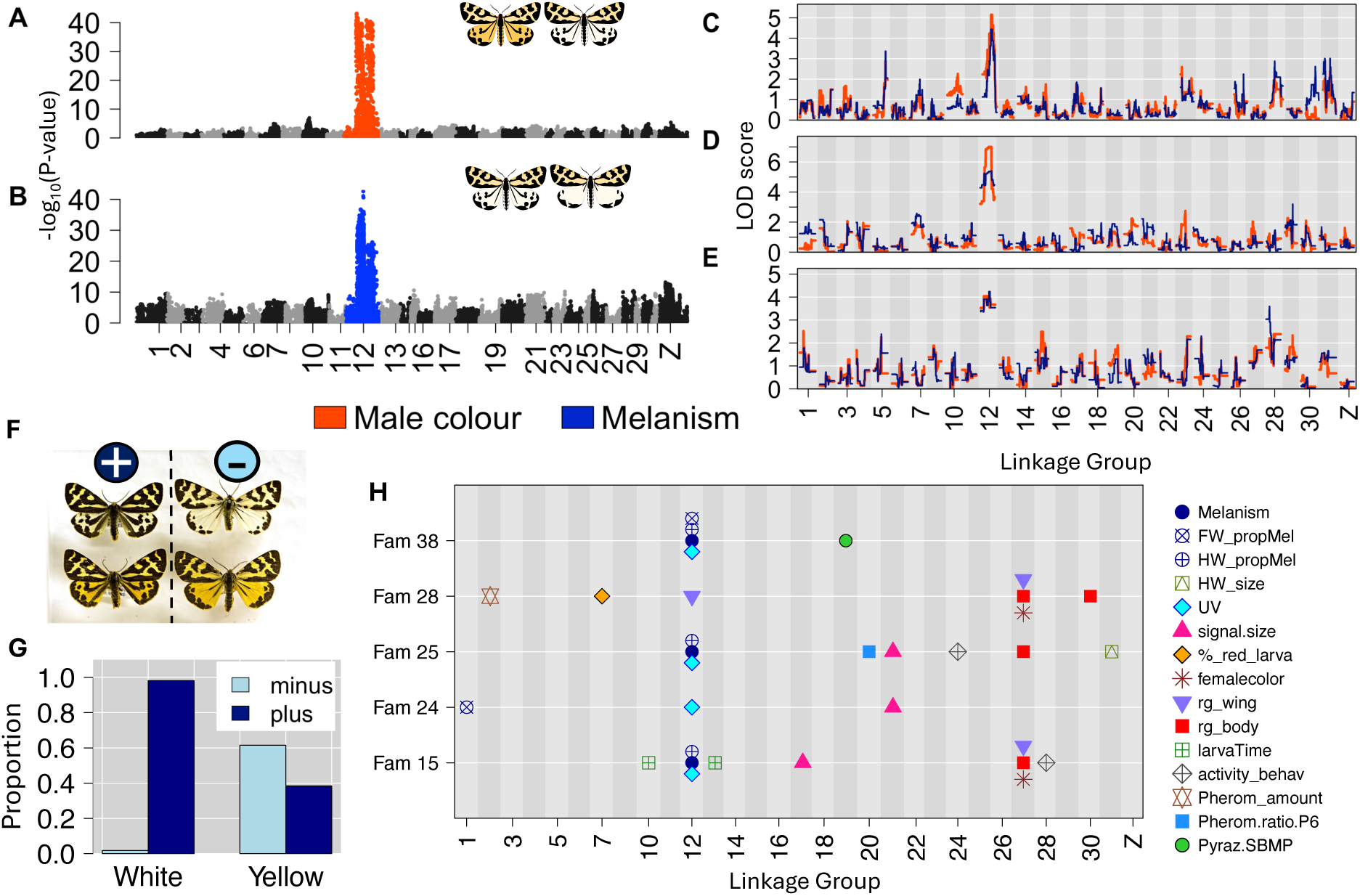
Genome wide association of male hindwing colour (A) and melanism type (B) and quantitative trait loci (QTL) mapping of both traits in three different families (family 15, 25, 38, C-E). Both were analysed as binary traits: male hindwing colour can be white or yellow, melanism types are plus and minus (F). They show a strong association in our data set (G), which corresponds to the underlying genetic bases, which shows almost identical associations with linkage group 12 (A-E). Apart from melanism (classified as plus/minus, proportion of melanisation on forewing and hindwing (‘FW_propMel’, ‘HW_propMel’), and UV brightness, there were no other traits that showed a strong association with the male colour locus. QTL mapping of different traits showed only one significant QTL for female hindwing colour (redness = ‘rg_wing’) in one of the families (H). However, there were significant associations between female colour measures and LG 27 detected in several families and with larval signal size and LG 21. See Figure 3 for more details and main text for descriptions of the different traits tested.

Six families from the first generation of 2021 (generation 24 of the laboratory population) that showed segregation of the male colour allele (Table 1) were selected for quantitative trait loci (QTL) analysis. Parents and grandparents of these families were also included in the sequencing. They were kept in the same way as the main stock population described above, apart from keeping larvae separately instead of in groups from the third instar onwards. This allowed us to measure larval signal sizes and assign these to other trait measurements (the signal disappears when an individual pupates). Signal size of larvae at the last instar was assessed by counting the number of body segments with orange hair. Signal size records in the laboratory population were only available for a subset of individuals.

**Table 1.** Sample sizes of the different families sequenced, the number of yellow and white males and plus and minus males within each family.

| *Family ID* | *N* | *Males* | *Females* | *white* | *yellow* | *Melanised plus* | *Melanised minus* |
| --- | --- | --- | --- | --- | --- | --- | --- |
| 15 | 135 | 63 | 72 | 36 | 27 | 37 | 24 |
| 24 | 125 | 63 | 61 | 53 | 10 | 61 | 1 |
| 25 | 100 | 52 | 48 | 40 | 11 | 37 | 13 |
| 28 | 155 | 71 | 84 | 55 | 16 | 70 | 1 |
| 38 | 73 | 38 | 35 | 28 | 10 | 27 | 10 |
| 49 | 43 | 22 | 21 | 17 | 5 | 19 | 3 |

### Phenotypes

The phenotypes included in the QTL analysis consisted of those that are measured in all individuals in the laboratory population, larval time (days from hatching to pupation), pupa weight, and melanism type. In addition, activity of adult males, amount and composition of pheromones (females only) and defence fluids (both sexes,) female wing and body colouration, and male wing UV reflectance were measured. For sample sizes see Table 2 (QTL mapping) and Table S1 (used for phenotypic analyses including those where genotyping failed).

**Table 2.** Samples sizes for traits used in the quantitative trait loci (QTL) analysis in the different families. Some traits were measured in both sexes, other traits were specific to one sex. Pyrazines: concentration of defensive compounds in defensive neck fluids; Signal Size: size of the warning signal in larvae quantified by counting the number of segments with orange hair; Prop.Col Larva: proportion of orange of larvae quantified from images; rg_larvaePatch: colour of the orange patch; Forewing/Hindwing_propMel: proportion of the wing that is melanised; Melanism: melanism type classified as plus or minus (see Figure 1F); UV brightness: UV brightness of male hindwings; Activity Behaviour: minutes active during a 50-minute behaviour assay; Female Colour: colour of female hindwings measured by visual matching (see Figure 3J); rg_body and rg_wing: red colour quantified from images.

|  | **Family** |  |  |  |  |  |  |
| --- | --- | --- | --- | --- | --- | --- | --- |
| **Trait** | **15** | **24** | **25** | **28** | **38** | **49** | **Total** |
| Larva Time | 135 | 121 | 99 | 153 | 73 | 43 | 624 |
| Pupa weight | 135 | 121 | 99 | 154 | 73 | 43 | 625 |
| Pyrazines | 42 | 43 | 36 | 43 | 40 | 22 | 226 |
| Signal Size | 135 | 125 | 100 | 155 | 73 | 43 | 631 |
| propCol.larva | 39 | 43 | 36 | 51 | 15 | 16 | 200 |
| rg_larvaePatch | 49 | 55 | 47 | 71 | 27 | 20 | 269 |
| **male traits** |  |  |  |  |  |  |  |
| Forewing Size | 59 | 64 | 49 | 70 | 37 | 19 | 298 |
| Hindwing Size | 59 | 63 | 47 | 66 | 37 | 20 | 292 |
| Forewing_propMel | 59 | 61 | 39 | 59 | 36 | 16 | 270 |
| Hindwing_propMel | 58 | 62 | 47 | 65 | 36 | 20 | 288 |
| Male Colour | 63 | 63 | 51 | 71 | 38 | 22 | 308 |
| Melanism | 61 | 62 | 50 | 71 | 37 | 22 | 303 |
| UV brightness | 60 | 64 | 50 | 69 | 37 | 17 | 297 |
| Activity Behaviour | 61 | 63 | 47 | 63 | 37 | 18 | 289 |
| **female traits** |  |  |  |  |  |  |  |
| Female Colour | 71 | 57 | 44 | 80 | 34 | 20 | 306 |
| rg_wing | 71 | 59 | 45 | 81 | 34 | 20 | 310 |
| rg_body | 72 | 55 | 46 | 80 | 33 | 20 | 306 |
| Pheromones | 40 | 39 | 22 | 58 | 16 | 11 | 186 |

### Melanisation, female colour and larval signal size

Male hindwing melanism type was classified as plus and minus based on presence or absence of a dark patch (Figure 1) and compared with proportion of melanised fore- and hindwings (Figure 2) quantified from images (see below). Colouration of the female hindwings was quantified by visual mapping against a colour scale (Figure 3) and extracted from photos. We measured the size of the larval signal size by counting the number of body segments with orange hair and quantified the colour as well as the proportional signal size based on image analysis.

**Figure 2.**
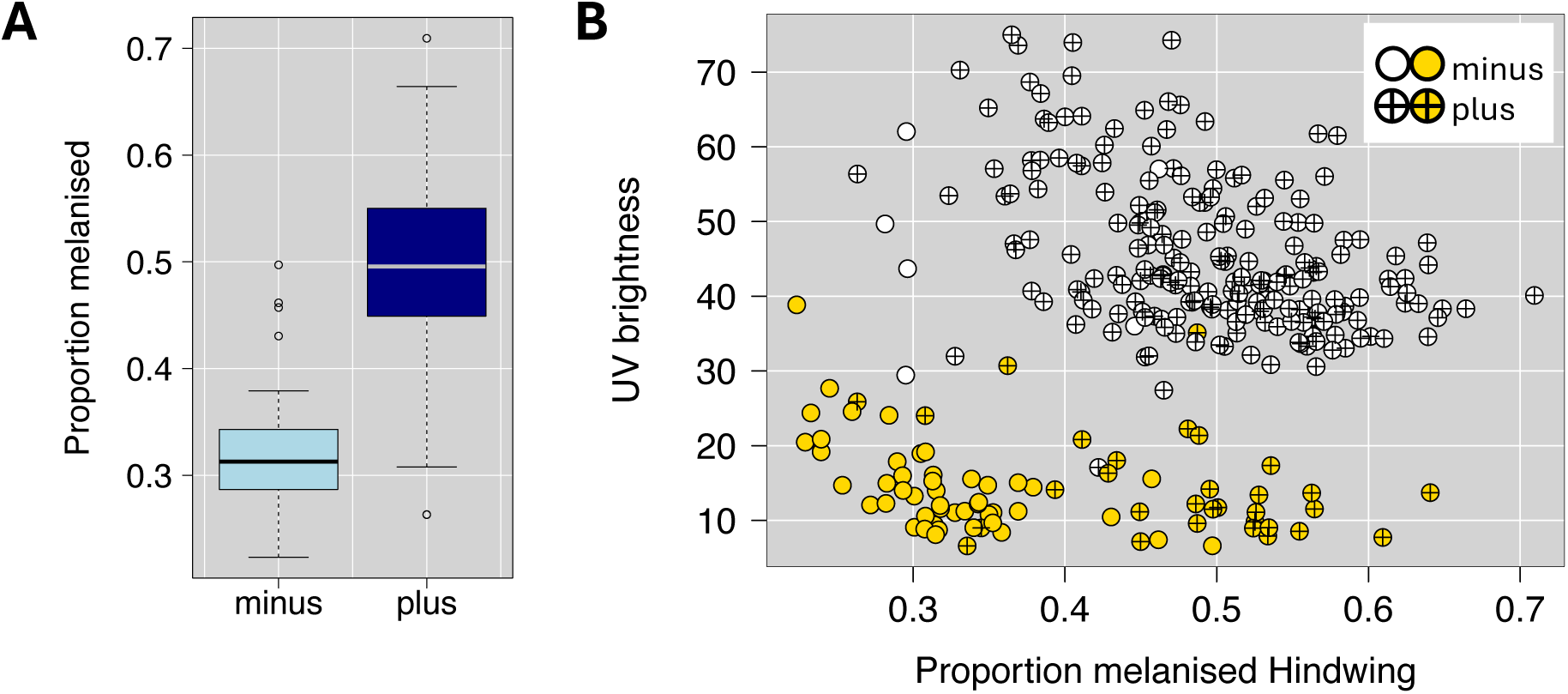
A: Relationship between melanism type (plus/minus, see Figure 1F) and the proportion of melanised hindwings quantified from images. B: Relationship between UV brightness of hindwings and the melanised proportion. Colours indicate male hindwing colour (yellow or white). Symbols refer to the melanism type.

**Figure 3.**
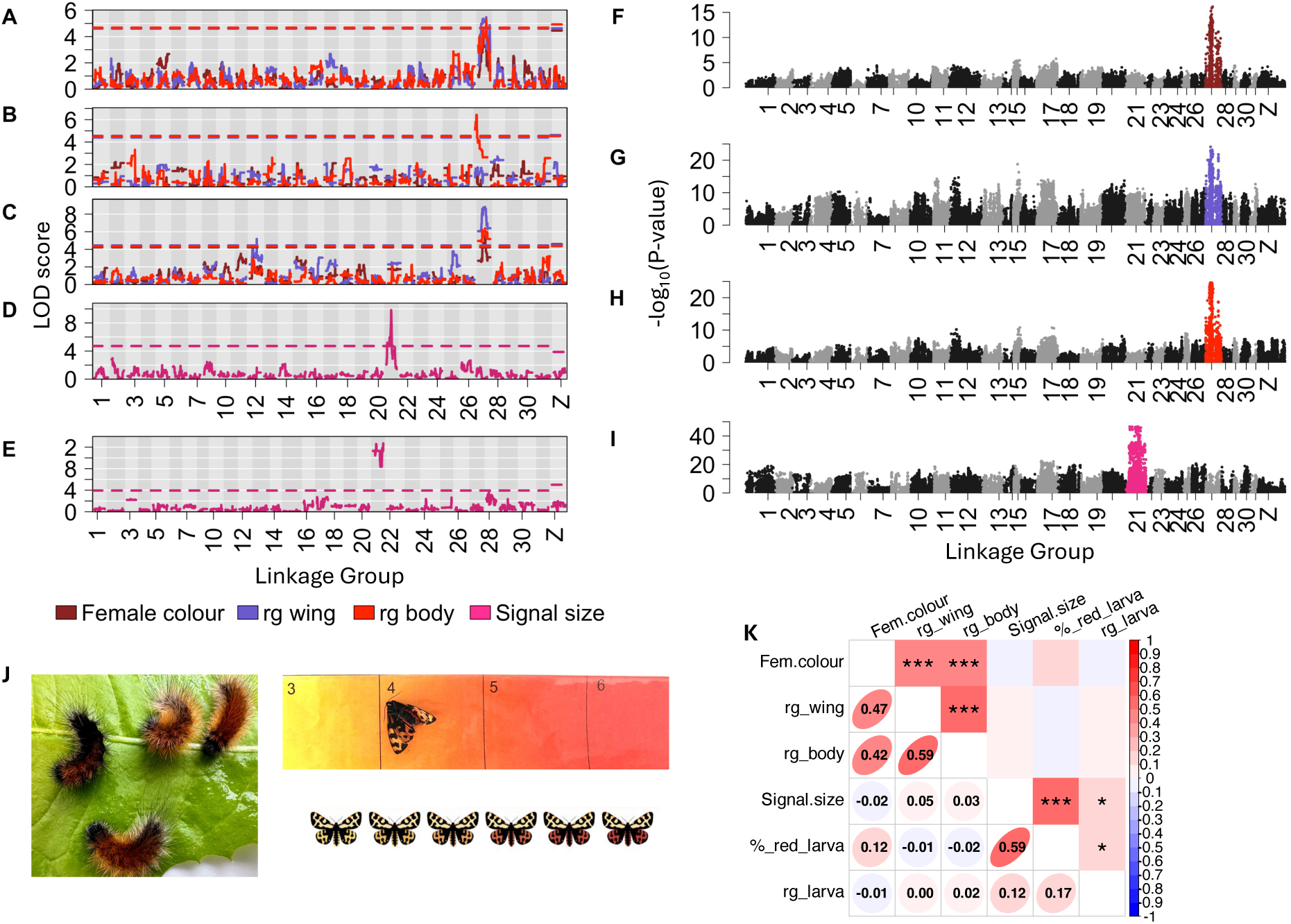
Quantitative trait loci analysis of female colour traits (A-C) and larval signal size (D, E) in different families (A: family 15, B: family 25, C: family 28, D: family 24, E: family 25). Hind wing colour was measured by visual matching against a colour scale (see J), here ‘Female colour’ (redness), and was also quantified from images (‘rg_wing’). Body colour was quantified from images (‘rg_body’). Larval signal size was measured by counting the number of segments with orange hair (J). Genome wide association analyses including all sequenced individuals confirmed the presence of large effect loci for female colour and larval signal size that were detected in the QTL analysis (I-J). Female colour traits are correlated (K), these includes the different measurements for hindwing colour as well as body and hindwing colour. The different measurements for larval signal size (‘Signal.size’ = number of segments with orange hair; ‘%_red_larva’ = from images) are correlated. The correlation between size of the orange patch and its colour (‘rg_larva’ measured from images) was not significant (K). Female and larvae colour traits show no significant correlation (K), which is consistent with the presence of QTLs on different linkage groups.

### Image analysis

Visible light and UV photographs were taken of late-stage larvae, adult wings and adult bodies under standard lighting conditions. For larvae, the size of the red patch was calculated as the number of pixels in the red region divided by the number of pixels in the whole body region. The proportion of melanised to non-melanised regions of the forewings and hindwings were calculated for the male moths. UV brightness was calculated for the non-melanised regions of the hindwings. For the female moths, redness of the coloured regions of the hindwings and the coloured areas of the body were measured. In all cases, redness was calculated as a ratio of red to green using RGB values in the formula (R-G)/(R+G). For all details see Appendix in Supporting Information.

### Activity trials

We tested the activity levels of adult males during 45-minute trials. Only males were tested since they are the more active sex. Prior to the trials, males were placed in cool boxes with ice for 30 minutes, so that all moths began the trial in a cooled state. They were then moved to individual plastic boxes, placed on their backs and observed for 45 minutes. During this time, we recorded: time until the moth flipped from its back to its front, number of minutes during which there was any antenna movement, walking, fluttering of the wings or flying, sum (in minutes) of all activities observed, and the number of minutes when any forms of activity was observed. Ambient temperature was recorded in the cool box, and in the room at the beginning and end of each trial.

For the QTL analysis, behaviour measurements were corrected for weather (factor: cloudy, half cloudy, sunny/half cloudy, sunny), and temperature using the residuals of linear models that included these confounding factors. To ensure normality, residuals were square-root transformed after adding the minimum residual value to obtain only positive values. Linear models and transformations for behaviour measurements, as well as for pheromones and pyrazines (see below), were performed in R 4.4.0 (R Core Team 2024).

### Chemical defence fluid

Thoraic fluid was extracted after eclosion by squeezing the moths’ thorax with tweezers and collected in a 10ul glass capillary. We measured concentration of SBMP (2-sec-butyl-3-methoxypyrazine) and IBMP (2-isobutyl-3-methoxypyrazine), the main defence compounds in the defensive neck fluids released by adults following the methods of Cai et al. (2007) as described in Burdfield-Steel et al. (2018). The amounts of IBMP, SBMP, their combined amounts and their ratio were log-transformed before analysis. More details in the Appendix, supporting information.

### Female sex pheromones

Female sex pheromones were collected when females were 1-2 days old. We measured the amount of two compounds, P3 and P6. Absolute amounts (in ng) of each compound were calculated relative to the 200 ng of internal standard. Finally, the ratio of the second major compound (P6) was calculated over the amount of the major compound (P3). Measurements of the total pheromone amount as well as P6.P3 ratio were log-transformed. More details in the Appendix.

### Sequencing and association analysis

#### Sequencing

Whole-genome haplotag sequencing was performed on 657 samples as described in Pal et al. (2025). Haplotagging is a linked-read sequencing technique that uses molecular barcodes in Illumina short-read sequencing that allowed downstream analysis to preserve haplotype linkage information (Meier et al. 2021). Following library preparation, samples were sequenced at ∼0.5× coverage with 150 bp paired-end reads. Immediately following sequencing, raw sequence data were demultiplexed and converted into standard FASTQ format supplemented with BX barcode tags. BX tags were retained in the read header for BX-aware duplicate removal and to inform read-aware genotype imputation downstream.

#### Sequence analysis

FASTQ reads were mapped to the white reference (Yen et al. 2020) using bwa-mem v7.12 (Li and Durbin 2009). BAM files were sorted and indexed using samtools version 1.10 (Li et al. 2009). PCR duplicates were identified and removed with MarkDuplicates from Picard (http://broadinstitute.github.io/picard/), version 2.9.2 with options READ_ONE_BARCODE_TAG=BX READ_TWO_BARCODE_TAG=BX. We called variants with bcftools v.1.9 (Li 2011) mpileup and call (--max-depth 10000 -q 30 -Q 20) using the multiallelic calling mode (Danecek et al. 2016). Resulting VCF-files of individual scaffolds were concatenated using bcftools. We filtered VCFs using vcftools v.1.14 (Danecek et al. 2011) where we only kept a set of the most reliable biallelic variants (--maf 0.01 --min-alleles 2 --minDP 1 --minGQ 4.2 min-meanDP 0.5 –minQ 100 max-missing 0.4), resulting in 325,944 variants.

#### Genotype imputation and phasing

Our sequencing strategy involved sequencing a large number of samples (657) at relatively low coverage, which resulted in a high proportion of missing data. However, because haplotagging provided information on linked reads, and the individuals were closely related—resulting in large haploblocks (high numbers of linked variants)—it was possible to reliably impute missing genotypes. We used STITCH (Davies et al. 2016) using the following parameters: --buffer=25000 --K=10 --method=diploid --nGen=30 --nCores=10 --readAware=TRUE --keepInterimFiles=FALSE --shuffle_bin_radius=500 --expRate=5 --iSizeUpperLimit=500000 --keepSampleReadsInRAM=FALSE -- inputBundleBlockSize=50000 --use_bx_tag=TRUE. We applied these parameters to genome wide windows of 500 kb. The resulting VCFs were merged using bcftools.

#### Linkage Map Construction

We constructed linkage maps using Lep-MAP3 (Rastas 2017) with the filtered VCF file as input. We first called parental genotypes using ‘ParentCall2’ and filtered the data based on segregation distortion with ‘Filter2’ (with removeNonInformative=1 and setting a more stringent threshold: dataTolerance=0.0001 (default=0.01)). We used a previous assignment of scaffolds to linkage groups (LG) based on Brien et al. (2022) after correcting erroneous assignments (see Appendix and Figure S1, S2). We ordered markers of each LG using ‘OrderMarker2’ with recombination2=0 (which accounts for no recombination in females) grandparentPhase=1 outputPhasedData=1 hyperPhaser=1 phasingIterations=12 usePhysical=1. We ran the ordering step six times for each LG and kept the run with the highest likelihood. We repeated this step in the same way without grandparental phasing (grandparentPhase=0) in families where grandparental information was missing. Isolated markers at the end of LGs, which could not be reliably placed and led to long gaps, were manually removed. The resulting linkage maps consisted of 31 LGs ranging in size from 74.1-99.0 cM with a total length of 2683.53 cM and 9227 unique positions (grandparental phased), and 68.9-97.9 cM with a total length of 2411.10 cM and 8133 unique positions without grandparental phasing. Phased data were converted for quantitative trait loci (QTL) mapping using Lep-MAP’s map2genotypes.awk script. In addition to the filtered VCF file, we constructed linkage maps using a phased VCF with missing genotypes imputed by STITCH (see Appendix, Supporting Information).

#### Sex Chromosome Identification

The species has a ZW sex determination system (Yen et al. 2020), but the sex chromosomes among the linkage groups or scaffolds had not been identified before. Knowledge of the sex chromosome is crucial for QTL analysis since it needs special statistical treatment and is essential for analysing sexually dimorphic traits. We identified the Z-chromosome based on relative sequencing depth. LG 8 showed only half the read depth in females compared to other LGs but no differences in males (Figure S3). We confirmed a highly significant association between this LG and sex (Figure S4) using a genome wide association analysis (GWAS). Additionally, 99.8 % of the markers identified as sex-linked by Lep-MAP3 were assigned to LG 8. More details in the Appendix.

#### QTL mapping

We used rQTL v. 1.66 (Broman et al. 2003) for QTL mapping within the R framework (R version 4.4.0 (R Core Team 2024)). Since the crossing to produce the individuals in our data set did not involve two distinct genetic groups, it was not possible to combine the different families (i.e. phasing is not comparable between families: for example, alleles that are inherited from the maternal grandmother are not expected to be the same in the different families). We analysed each family separately as a 4-way QTL cross using Haley-Knott regression. Family 49 was too small for a QTL analysis (Table 1). Male colour and melanism were analysed as binary traits. For traits that were not specific to one sex, we included sex as a covariate (pupa weight, larval signal size, larval time, pyrazines (IBMP, SBMP, and their ratio). Significance thresholds were obtained by running 10,000 permutations. Confidence intervals for the position of a QTL were inferred using the lod_int function in rQTL.

#### Genome wide association analysis

We performed a genome-wide association study (GWAS) using ANGSD (Korneliussen et al. 2014), which accounts for genotype uncertainty. As input, we used the VCF file with missing genotypes imputed by STITCH (see above), keeping only variants with an INFO score greater than 0.5. ANGSD was run with the parameters - doMaf 4 -vcf-gp -doAsso 2. Family was included as a covariate. Male colour, melanism, and sex were treated as binary traits, while female colour measures and larval signal size were analysed as continuous traits. A recessive model was applied for male colour (-model 3).

#### Candidate genes

To test whether detected associations included regions with genes known for colour patterns in Lepidoptera, we ran a protein BLASTP v2.16.0 search to find homologous regions in the *A. plantaginis* WW reference genome corresponding to the candidate genes *optix, cortex, wnta, domeless, vvl* (Meier et al. 2021; Van Belleghem et al. 2021) shown to be involved in wing colour and patterning in Lepidoptera or *Drosophila* (*ebony, tan*) (Massey and Wittkopp 2016).

#### Comparison of yellow and white morphs in pedigree population

We used the extensive data set of the whole laboratory population collected over 25 generation to test for differences in larval time, larval signal size, and pupa weight between the yellow and white morphs. We used linear mixed model including year (as factor), generation, month (factor), inbreeding coefficient, and melanisation type, as fixed effects and family as random effect to account for non-independence of full-sibs using the Rpackage ‘lme4’ (Bates et al. 2015). Significance was assessed after correcting for the mentioned fixed factors. Inbreeding coefficients were calculated from the pedigree using the Rpackage ‘pedigree’ (Coster 2022). Additionally, we tested for effects of melanisation after correcting for male colour effects and within each of the colour morphs separately. We also tested for interactions between male colour and melanism. Larval times of overwintering individuals were excluded. All statistical analyses were conducted in R version 4.4.0 (R Core Team 2024).

#### Effects of male colour and melanism within sequenced individuals

We examined the effects of 1) male colour genotype and melanism (+/- type) and 2) colour genotype and melanism genotype on other traits among the subset of sequenced individuals using linear mixed models in the R package ‘lme4’. For these models, we selected the SNP genotypes that showed the strongest associations in the GWAS analysis: WW_tarseq_419_arrow.6870348 for male colour and WW_tarseq_540_arrow.17610360 for melanism type using the VCF with missing genotypes imputed by STITCH. Additionally, sex was included as a fixed factor in the second model (2), while family was treated as a random effect. Using genotypes provided the advantage of an increased sample size, as it allowed us to include females in 2), for which we lack phenotypic data for these male-specific traits. Additionally, male colour and melanism are strongly correlated in our dataset (with almost no white/minus individuals), making it difficult to disentangle their potential effects. In contrast, genotypes provided more informative data, improving our ability to analyse these associations

We tested for associations between activity and male colour using linear mixed models (Rpackage ‘lme4’ (Bates et al. 2015)). We included weather, temperature and male colour as fixed effect and family as random effect. Since it is known that melanisation can have a substantial impact on thermoregulation and the temperatures during the behaviour assay were relatively high (26-34 °C) we included wing melanisation as well. Distribution of residuals was examined to evaluate validity of models.

#### Associations between other traits

Many of the traits studied here, along with their genetic correlations and associations with warning colour variation, have been explored in a previous study (Koch et al. 2024). In this study, we also examined additional traits, including pyrazines and pheromones. To test for associations between traits, we used linear mixed models with family as a random factor, as described above. Specifically, we tested the effects of pyrazine concentrations on pupa weight, larval time, melanisation, signal size, and female colour. For pupa weight and larval time, sex was included as a fixed factor, and we also tested for interactions between pyrazine concentrations and sex. Additionally, we examined differences in pyrazine concentrations between sexes using linear mixed models with family as a random effect. Finally, we tested for associations between the total amount of pheromones, pheromone composition (P6-P3 ratio), and pupa weight, larval time, female colour, signal size, and pyrazines. Sample sizes for all traits are provided in Table S1.

## Results

### Genetic basis of phenotypic variation

We explored the genetic basis of different traits in *A. plantaginis* and were specifically interested in whether any showed associations with the male colour region. We sequenced 657 individuals from generations 24 and 23 of the laboratory population. We generated a linkage map and used QTL mapping to find associations between genomic and phenotypic variation in a wide range of different traits including size, developmental time, colouration in larvae and females, male activity, pheromone amount and composition and composition of defensive fluids. Additionally, we used a GWAS analysis to confirm the presence of QTLs detected in several families independently.

### Linkage maps

Our linkage map improved the genome assembly. We could identify the Z-chromosome (LG 8, see Figure S3) and assigned parts of the scaffold WW_540_tarseq to the LG containing the male colour locus (see Appendix, Figure S1, S2). We conducted the QTL analyses with two data sets. Set 1) consisting of a filtered VCF and 2) a VCF with missing genotypes imputed and phased by STITCH, to explore whether genotype imputation and using the phase information of the haplotagging data during linkage map construction can increase the power to detect QTLs (see Appendix). The results were very consistent (see Table S2), with the exception of a QTL for larval signal size on LG 21 in family 25, which we could only detect with the STITCH imputed data set, but which was found in another family using the filtered VCF set and confirmed by a GWAS. We further found that traits with clear signals in some families but not reaching the significance threshold in others (melanism, UV brightness in family 15, signal size in family 25), were significant using the STITCH imputed data set. However, all linkage maps obtained from the STITCH imputed data were severely inflated with LG length between 141 and 316 cM.

### Quantitative trait locus mapping

For most of the measured traits we could not detect QTLs that were consistent across several families. They were often detected in only one of the families or differed in locations between families. Exceptions are melanism and UV light reflectance that consistently mapped to LG 12 at the location of the male colour locus. Our different measurements of melanism, melanin type (plus/minus) and proportion of melanised hindwings, showed consistent QTLs (Figure 1). In one family we also found a significant QTL for forewing melanisation on LG 12. Furthermore, we found strong evidence for a QTL for female hindwing colour on LG 27 with consistent results in three families and for larval signal size on LG 21 in two families, more details in respective sections below.

### Associations between male colour, wing melanism, and other traits

We see an increase of the melanised form over time in the laboratory population, especially within the white males (Figure S5). In the first generation, among the founder individuals, plus and minus melanisation occurred at similar proportion in yellows and whites, but over the following generations, the plus type reached almost 100 % in white males while staying at around 40 % in yellows. Given the known costs of melanin synthesis in insects (True 2003; Talloen et al. 2004), we therefore accounted for melanisation when testing male colour effects. We detected a significant association between male colour and larval time with yellow individuals developing slightly faster (−0.26 ± 0.11 days; P = 0.02). However, this effect could only be detected when analysing the total data set comprising of data from 2013 to 2021 and covering 25 generations and disappeared when analysing the last years (2018-2021) separately (Table 3). This may suggest that strong lab selection, indicated by the dramatic decrease in developmental time (Figure S5), eroded some allelic variation responsible for developmental time that was linked to the colour locus in the early generations. Associations with other traits were not significant (Table 3). We found no effect of melanism type on larval time, pupa weight and larval signal size, no significant differences in any trait between melanisation types within colour morphs, and no significant interaction between male colour and melanisation (Table 3).

**Table 3.**
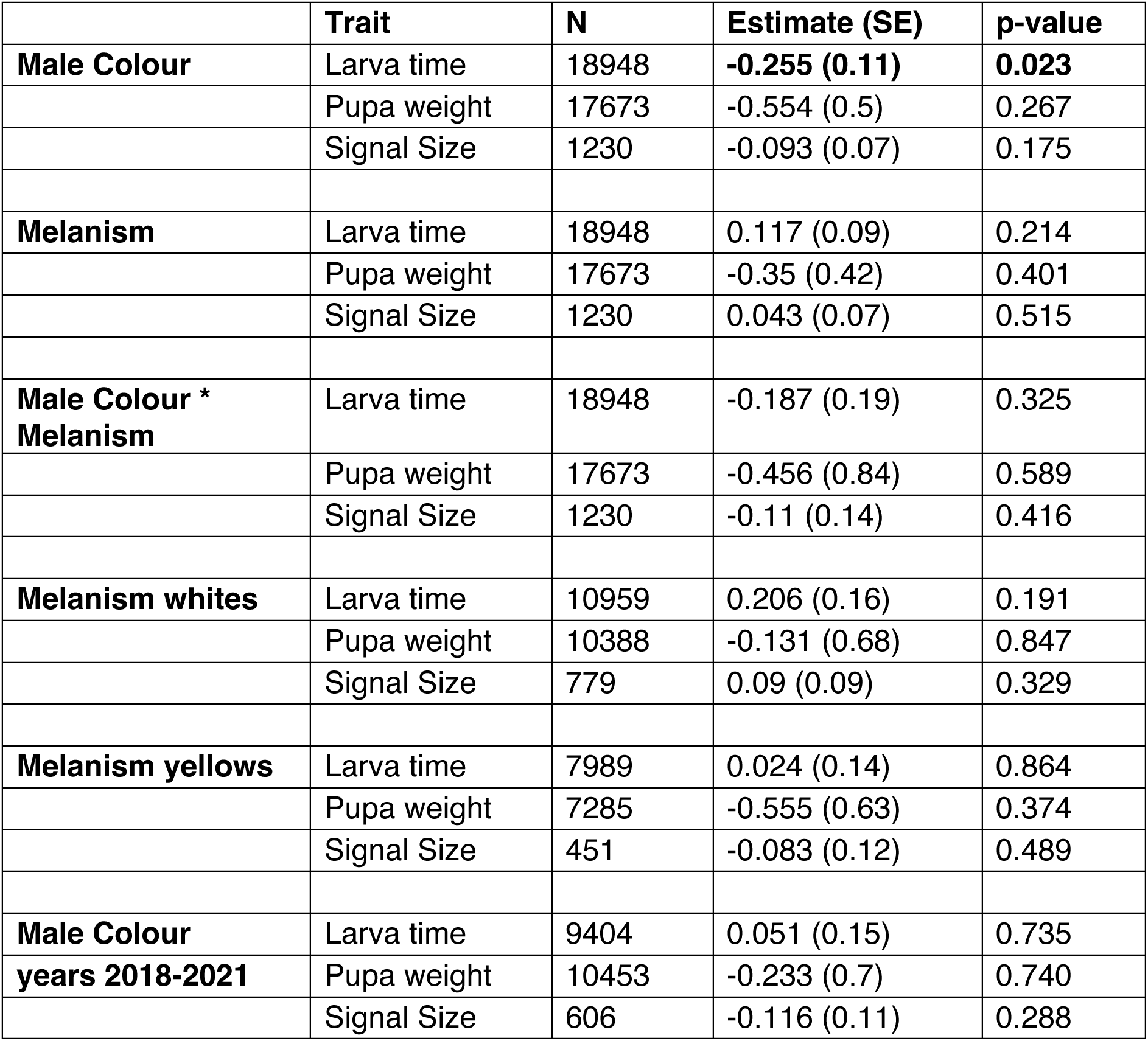
Effects of male colour (white/yellow) and melanism type (minus/plus) on different traits estimated using linear mixed models. Significant estimates are shown in bold. We tested male colour and melanism effects in the complete laboratory population including years from 2013-2021 and spanning 25 generations, melanism effects within each colour morph (complete population), and male colour effects in the last four years.

The sequenced individuals were from the same laboratory population. Accordingly, male colour and melanism are strongly associated in our data set (see Figure 1G), with almost all whites showing the melanised form, whereas yellows consisted of both melanin types. Our genetic analysis confirmed that a major effect allele for melanism is closely linked to the *valkea* gene: In three families we detected a QTL for melanism type on the same LG and showing the same association pattern as male colour (Figure 1C-E, Table 4, S2). Due to a lack of variation in melanisation, QTL mapping in the other families (families 24, 28, 49) was not possible (Table 1). A GWAS rediscovered the known male colour locus on LG 12 (Figure 1A) and showed almost identical associations between genomic regions with male colour and melanism (Figure 1B). However, there are subtle differences and the fact that all possible combination of hindwing colour and melanism occur (i.e. yellow plus and white minus, Figure 1F) rules out a pleiotropic effect of *valkea* on melanism. In the GWAS, we observe the strongest association with melanism for SNPs on scaffold WW_540_tarseq, whereas male colour shows almost equally strong signals to WW_540_tarseq and WW_419_tarseq (see Figure 1A showing the broad peak for male colour versus the narrower peak for melanism in Figure 1B). An additional GWAS with yellow males only confirmed the presence of a large effect allele on LG 12 (see Figure S6), which is independent of *valkea.* The top SNPs (-log_10_(p-value) > 43, see Figure S6) occurred on WW_540_tarseq:17997943-18721005, which contains the *cortex* locus (Table S3) that is well known for its role in Lepidoptera wing melanisation and pattern (Nadeau et al. 2016; van’t Hof et al. 2016; Wang et al. 2022; Zhang et al. 2024). However, among the yellow males the strongest association with melanism was found on a scaffold on the Z chromosome (scaffold WW_222_tarseq). This region showed a weak association in the GWAS with white males only as well indicating that variation in melanism may be controlled by several loci with different effect sizes.

**Table 4.** Results of quantitative trait (QTL) analyses in different families. Each family was analysed separately as a 4-way cross. LG= Linkage group. CI = Confidence interval. Behaviour: minutes of activity during a 45-minute behaviour assay; rg_body and rg_wing: red colour quantified from images; Signal Size: size of the warning signal in larvae quantified by counting the number of segments with orange hair; Prop.Col Larva: proportion of orange of larvae quantified from images. UV brightness: UV brightness of male hindwings; Melanism: melanism type (plus/minus, see Figure 1). For results obtained from STITCH imputed data see Table S2.

| Famil<br>y | Trait | LG | LG<br>positio<br>n [cM] | CI<br>lower | CI<br>high<br>er | LO<br>D | P-value | Scaffold | Position<br>on<br>Scaffold |
| --- | --- | --- | --- | --- | --- | --- | --- | --- | --- |
| 15 | Behaviour: wing fluttering | 28 | 34.5 | 22.9 | 84.6 | 5.3 | < 0.001 | WW_tarseq_490_ar<br>row | 294264<br>5 |
| 15 | Larva Time | 13 | 72.5 | 71.5 | 80.1 | 4.9 | 0.031 | WW_tarseq_553_ar<br>row | 763154 |
| 15 | rg_body | 27 | 62.0 | 8.4 | 79.6 | 5.3 | 0.010 | WW_tarseq_565_ar<br>row | 27994 |
| 15 | rg_wing | 27 | 61.6 | 14.5 | 90.4 | 5.5 | < 0.001 | WW_tarseq_479_ar<br>row | 188157 |
| 24 | Forewing_prop.<br>melanised | 1 | 5.4 | 0 | 92.8 | 6.1 | 0.021 | WW_tarseq_521_ar<br>row | 173096<br>42 |
| 24 | Signal Size | 21 | 33.8 | 33.0 | 35.7 | 9.8 | < 0.001 | WW_tarseq_510_ar<br>row | 116128<br>5 |
| 24 | UV brightness | 12 | 47.7 | 36.4 | 54.8 | 8.7 | < 0.001 | WW_tarseq_419_ar<br>row | 696883<br>6 |
| 25 | Behaviour:<br>sum_activities | 24 | 26.6 | 0 | 59.1 | 6.2 | < 0.001 | WW_tarseq_505_ar<br>row | 951645<br>2 |
| 25 | Hindwing_prop.<br>melanised | 12 | 33.4 | 0 | 84.6 | 6.0 | 0.006 | WW_tarseq_487_ar<br>row | 621435 |
| 25 | Hindwing size | 31 | 53.6 | 33.0 | 54.7 | 4.8 | 0.027 | WW_tarseq_466_ar<br>row | 321747<br>3 |
| 25 | Melanism | 12 | 67.2 | 0.0 | 84.6 | 5.4 | < 0.001 | WW_tarseq_540_ar<br>row | 147941<br>42 |
| 25 | rg_body | 27 | 10.8 | 0.0 | 14.3 | 6.4 | 0.003 | WW_tarseq_524_ar<br>row | 203138 |
| 25 | UV brightness | 12 | 47.2 | 25.8 | 66.2 | 6.7 | 0.004 | WW_tarseq_515_ar<br>row | 954443 |
| 28 | Female colour | 27 | 52.9 | 32.5 | 59.6 | 5.5 | < 0.001 | WW_tarseq_608_ar<br>row | 36846 |
| 28 | Prop.Col Larva | 7 | 75.8 | 72.5 | 77.4 | 4.7 | 0.031 | WW_tarseq_489_ar<br>row | 112532<br>59 |
| 28 | rg_body | 27 | 53.1 | 0.0 | 92.8 | 6.4 | < 0.001 | WW_tarseq_565_ar<br>row | 97414 |
| 28 | rg_wing | 12 | 53.7 | 19.6 | 54.0 | 5.2 | < 0.001 | WW_tarseq_515_ar<br>row | 953622 |
| 28 | rg_wing | 27 | 52.9 | 27.7 | 63.4 | 8.8 | <<br>0.00<br>1 | WW_tarseq_608_ar<br>row | 36846 |
| 28 | Tot_phero_amo<br>unt | 2 | 83.2 | 56.4 | 89.8 | 7.0 | 0.00<br>2 | WW_tarseq_508_ar<br>row | 458401<br>1 |
| 38 | Forewing_prop.<br>melanised | 12 | 75.0 | 60.2 | 75.9 | 5.3 | 0.02<br>3 | WW_tarseq_540_ar<br>row | 214946<br>36 |

### Hindwing colour and melanisation influence UV brightness, but show no association with other phenotypes

In addition to the plus-minus classification for melanin, the proportion of melanisation of hind- and forewings was also quantified from images. We confirmed that the plus type had a significantly higher proportion of hindwing melanisation (t-test, t = −18.583, df = 93.031, p-value < 2.2e-16, Figure 2A). Hind- and forewing melanisation were highly correlated (correlation ± standard error (SE) = 0.83 ± 0.03, P < 2.2e-16). We found that male colour and hindwing melanisation significantly influenced UV brightness (Figure 2B). UV brightness was significantly lower in the yellow morph (estimate ± SE = −41.51 ± 4.4, P < 2.2e-16) and decreased with increasing melanisation (−35.13 ± 7.3, P = 2.12e-06). Both traits showed a significant interactive effect (20.79 ± 10.1, P = 0.040) indicating that the effect of melanisation leads to a stronger UV brightness decrease in the white morph compared to the yellow morph (Figure 2B). We could not detect any significant effect or significant interaction on any other trait when we tested for associations with the male colour genotype and melanism (Table S4).

### Behaviour is affected by melanisation but not male colour

We found that activity was primarily dependent on weather and temperature. Temperature during the activity assay was consistently high ranging between 26 and 34 °C. Activity decreased in sunnier conditions and with increases in temperature (Figure S7). We could not detect an effect of male colour on activity, but found that a higher proportion of melanised hindwings resulted in reduced activity and detected a significant interaction between hindwing melanisation and temperature (Table S5). Weakly melanised males seem to be more active at lower temperatures compared to strongly melanised males and showed a stronger reduction in activity in response to temperature increases whereas melanised ones remained relatively inactive across the temperature range tested (Figure S8). We found no effect for forewing melanisation (Table S5), possibly due to smaller variation in melanisation (coefficient of variation hindwings = 0.021; forewing = 0.010).

### Female and larval colour: Consistent results for major effect alleles

In contrast to the discrete male colour morphs, females show continuous variation in their hindwing colour, which we quantified in two different ways. The usual approach applied to all females in the laboratory population is to match them against a colour scale (Figure 3J, see also Lindstedt et al. (2016), Nokelainen et al. (2022a)). In this study, we also used image analysis to obtain a more precise quantification for the ‘redness’ of female hindwings (‘rg_wing’) as well as female bodies (‘rg_body’). Similarly, larval signal size had been measured by counting segments with orange hair (Lindstedt et al. 2010). Here, we quantified the proportion of the orange patch and the redness of the larval signal (‘rg_larva’) from images.

### Correlation between measures of colour

Our two measures of female hindwing colour were significantly positively correlated (correlation [confidence interval]: 0.47 [0.38, 0.55], P < 2.2e-16) confirming that the visual matching approach captures meaningful variation (Figure 3K). Interestingly, both measures of female hindwing colour showed a positive correlation with body colour (female colour-rg_body: 0.42 [0.33, 0.51], P = 2.015e-15; rg_wing-rg_body: 0.59 [0.52, 0.66], P < 2.2e-16). The two measures of larva signal size were positively correlated (0.59 [0.49, 0.67]), whereas the correlation between signal size and signal colour was positive but smaller. While the correlations within larval and female colour traits were significantly positive, correlations between colour traits across different life stages (between larval and adult female colouration) were very weak and non-significant suggesting that these traits are largely independent.

### QTL analysis and GWAS

The measurements for female hindwing colour as well as body colour showed consistent QTLs in three families on LG 27 (Table 4, Table S2), which was also confirmed by a GWAS based on all samples (Figure 3F-H). For larval signal size we detected a significant QTL peak on LG 21 in two families (Figure 3D, E, I), which also showed a clear association in the GWAS. We found that *optix,* a gene well-known for its role in controlling red wing colour in Lepidoptera (Reed et al. 2011), was on LG 21, which showed a strong association to larva signal size. However, closer inspection (Figure S9) revealed that *optix* was outside the region of the highest association. None of the other candidate genes were on the LG associated with female colour.

### Chemical defences and pheromones and their associations to other traits

Associations between several of these traits and their genetic correlations in this specific population have been investigated before (Koch et al. 2024). Novel traits in this study included chemical defence compounds in the defensive neck fluids (pyrazines: IBMP, SBMP) and pheromones for which we tested associations with other traits.

The concentration of IBMP, SBMP, their combined concentration, and ratio did not differ between sexes (see Table S6), but males released significantly more neck fluids (log(volume) estimate ± SE = 0.638 ± 0.06, P < 2e-16). There was no association between neck fluid volume and pupa weight when accounting for sex (estimate ± SE = 8.80e-04 ± 1.1e-03, P = 0.42). When testing the relationship between fluid composition and other traits, we found that higher SBMP concentrations were associated with smaller pupa weight in both sexes (estimate ± SE = −5.33 ± 1.8, P = 0.004, Figure 4A). Relationships between larval time and pyrazines were less clear and differed between sexes (Table S6). SBMP was negatively correlated with larval time but showed a significant interaction with sex. The IBMP-to-SBMP ratio was positively correlated with larval time in both sexes (estimate ± SE = 0.92 ± 0.35, P = 0.009), while total pyrazine concentration showed no significant effect. No significant associations were found between single pyrazine compounds, their combined concentrations or their ratio and melanisation (fore- and hindwings), larval signal size, or female colour measurements for hindwings and bodies (Table S6).

**Figure 4.**
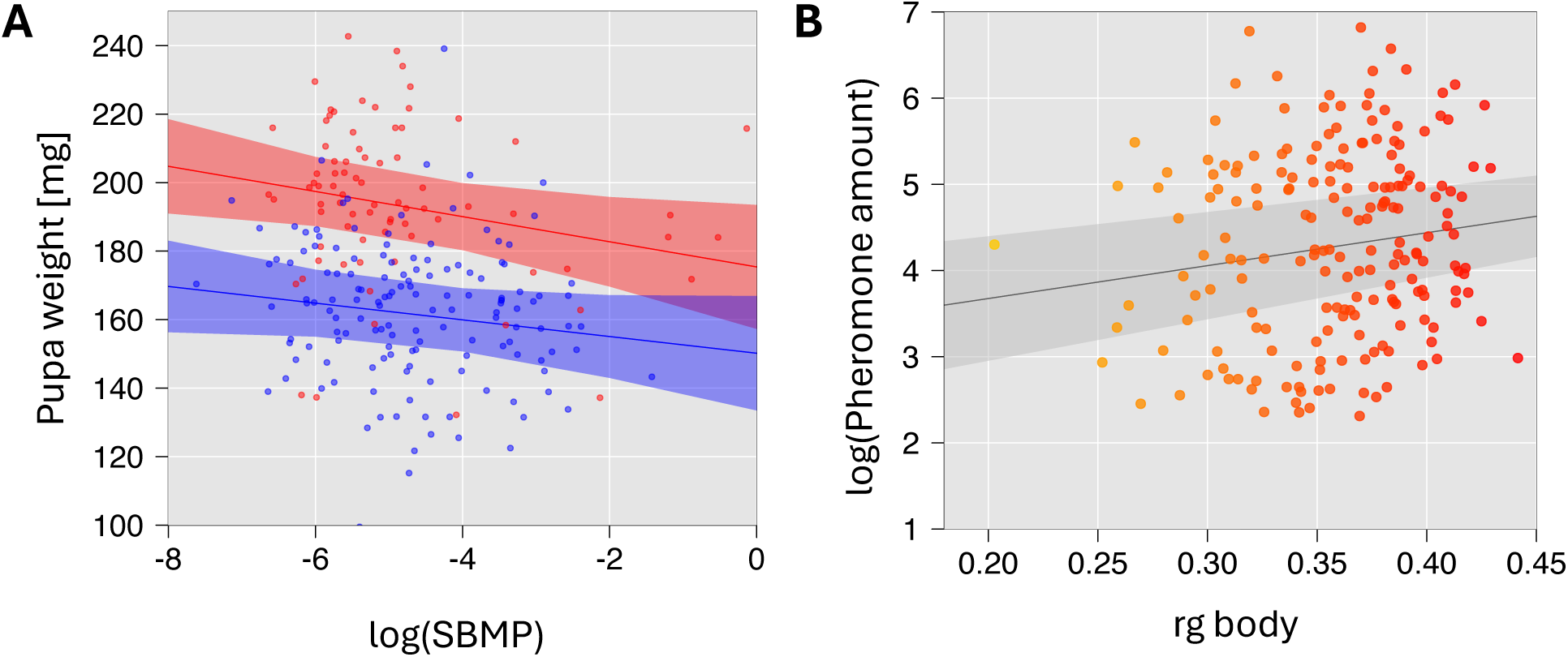
A: Association between the concentration of the SBMP (2-sec-butyl-3-methoxypyrazine), a major compound of the chemical defence in the defensive neck fluid, and pupa weight in males (blue) and females (red). Shown are predictions and confidence intervals from linear mixed models (see Method and Result for details) and observations. B: Relationship between total amount of pheromones and body colour (‘rg body’, ranging from yellow to red) in females. Model prediction and confidence interval are based on a linear mixed model.

We found a significant positive association between the total amount of pheromone and the redness of female hindwings (female colour: 0.10 ± 0.04, P = 0.03) and female body colour (rg_body: 5.51e-03 ± 2.68e-03, P =0.04; see also Figure 4B), but not with pheromone composition (P6.P3 ratio, see Table S6). We could not detect any associations between pheromone amount or composition (P6-P3 ratio) and pupa weight, larval time, larval signal size, and pyrazines (Table S6).

## Discussion

### Indirect selection by linkage between loci controlling male colour and hindwing melanism?

We aimed to determine if behavioural and life-history traits of *A. plantaginis* showed an association with the male colour locus. We found no evidence for an association between male colour and any of the measured traits, except for hindwing melanism and UV-light reflectance. The association with UV brightness is likely a direct consequence of hindwing colour and melanisation (white wings have higher UV reflectance). This may influence detectability by bird predators and can also be perceived by females (Henze et al. 2018), potentially contributing to mate choice. In contrast, the strong association between melanism and male colour is not due to pleiotropy but rather linkage between a locus controlling melanism and the previously identified male colour locus. Given the impact of melanism on multiple traits in insects (True 2003; Talloen et al. 2004; Ethier et al. 2015), this could have been a potential mechanism generating differences in other traits between colour morphs as well as maintaining their polymorphism by indirect selection. Melanin is associated with slower larval development (Ojala et al. 2007; Friman et al. 2009; Lindstedt et al. 2020; Koch et al. 2024), improved thermoregulation at low temperatures (Lindstedt et al. 2009; Nielsen and Mappes 2020) and increased pathogen resistance (Friman et al. 2009; Zhang et al. 2012) in this species. To assess potential associations between male colour and other traits, we therefore analysed it and melanism jointly, ensuring that any opposing linked effects did not obscure patterns. We found no evidence that either influenced developmental time, weight, or other colour traits. We detected slight differences in behaviour with darker individuals that are likely to heat up more quickly (Hegna et al. 2013) exhibiting lower activity than those with weaker melanisation at high temperatures. However, while the loci controlling male colour and melanism are linked, they are sufficiently distant to allow for recombination. Based on their positions on the scaffolds and the order of scaffolds on the linkage group, they are approximately 6.4 Mb apart, providing substantial opportunities for recombination in natural populations with a larger effective population size than the lab population. Consequently, indirect selection on male colour due to selection on melanism is likely to be minor.

### Maintenance of male colour polymorphism: No evidence for a complex phenotype

Maintaining the laboratory population over multiple generations may have reduced the initial genetic variation. However, both male colour morphs were maintained. If male colour had strong pleiotropic effects on other traits, we would have expected to detect them in our analysis. Inconsistencies between previous studies regarding trait differences between male colour morphs (eg. Nokelainen et al. 2012; Rojas et al. 2018 vs. Winters et al. 2021) may partly reflect differences in laboratory population history. Earlier generations were maintained under largely random mating and periodic influx from wild populations, whereas more recent matings have been structured to ensure adequate representation of specific genotypes in the population. (Gordon et al. 2018; Winters et al. 2021; De Pasqual et al. 2022; De Pasqual et al. 2024). Over time, this non-random mating between colour genotypes may have led to spurious associations between other traits and male hindwing colour within colour lineages. Additionally, it led to different levels of inbreeding between colour genotypes, which could have influenced reported trait differences and aligns with the occasionally observed superior performance of admixed heterozygous individuals (Gordon et al. 2018; Winters et al. 2021; De Pasqual et al. 2022). To avoid these confounding effects, we used families containing both white and yellow individuals, ensuring a shared genomic background across male colour morphs. Under this design, we were unable to identify any associated genomic loci for most of the measured traits or a phenotypic correlation with male colour, indicating that the male colour locus it is unlikely to be a major-effect locus for these traits. Given that previous research detected significant genetic variance in several of these traits (Lindstedt et al. 2009; Lindstedt et al. 2016; Koch et al. 2024), they likely have a polygenic basis with many small-effect loci contributing that we cannot detect with our sample size.

Taken together, the persistence of colour polymorphism in natural populations is most likely explained by spatial and temporal variation in selection acting directly on colour rather than associated traits. The effectiveness of warning signals is known to depend strongly on ecological context, including predator community composition (Nokelainen et al. 2014; Rönkä et al. 2020), presence of other aposematic species (Rönkä et al. 2018), and abundance of naïve juvenile birds during adult emergence (Mappes et al. 2014). While a conspicuous warning signal may confer an advantage in some environments, a more cryptic morph may experience higher survival in others.

This ecological perspective may also help explain the pronounced sexual dimorphism in the species. If predator warning was the main selective force, we might expect consistent warning colouration across sexes. Instead, females display red hindwings, the most effective warning signal (Rönkä et al. 2018; Rönkä et al. 2020; Gordon et al. 2021) and are largely sedentary. Males, in contrast, disperse actively and may benefit from reduced detectability during movement, suggesting that selection on male colour could be partly driven by crypsis rather than warning signal efficiency.

Additional ecological complexity likely arises from environmental variation. For example, light conditions can differentially alter predation risk between morphs (Nokelainen et al. 2022b), and there is evidence for predator driven positive frequency-dependent selection (Rönkä et al. 2020; Gordon et al. 2021). Combined with gene flow among populations (Galarza et al. 2014)), such spatially heterogeneous selection may facilitate the long-term maintenance of polymorphism. Theoretical models parameterized using data from *A. plantaginis* have already demonstrated that colour polymorphism can persist under positive frequency-dependent selection and migration between subpopulations (Gordon et al. 2015).

Finally, it is important to acknowledge that differences in fitness components like survival, dispersal ability, or mate-finding efficiency between morphs may exist, but are challenging to detect.

### Independent large effect loci for colour traits

In addition to the previously detected male colour locus, we identified major effect loci for female hindwing colour and larval signal size. These warning colour traits all map to different genomic regions, indicating that they can evolve independently. This aligns with previous findings (Nokelainen et al. 2022a) showing no association between male and female warning colour and suggests that there are no genetic trade-offs in warning signal efficiency across life stages or between sexes. Similarly, separate QTLs for female and larval colour and no overlap with QTLs for other traits aligns with a lack of genetic correlations between them as well as other traits reported previously (Koch et al. 2024). Interestingly, we also found no correlation between larval melanisation and male melanism, which fits to the detection of two different major QTLs located on separate linkage groups. Additionally, while larval melanisation was significantly associated with developmental time, male melanism showed no such correlation, further supporting the idea that these traits are regulated by distinct genetic and physiological mechanisms with different resource allocation trade-offs.

### Chemical defence: costs, sex differences, and lack of association with colour

Chemical defences (pyrazines) in this species are produced de novo (Burdfield-Steel et al. 2018) and become less effective when larvae are raised in protein-deprived environments (Burdfield-Steel et al. 2019; Ottocento et al. 2024), suggesting that they are costly to produce and require sufficient resources. We found that higher SBMP concentration in the defensive fluids, a major deterrent compound (Ottocento et al. 2023), was correlated with lower body weight and increased developmental time in males, indicating potential costs of producing this defence. Neck fluid volume did not correlate with weight showing that the total amount of pyrazine was indeed different between individuals of different sizes. While males produced a larger volume compared to females, the composition remained consistent across sexes. This pattern suggests that males invest more in chemical defence by producing a greater total amount of pyrazine, which may also explain the differing associations between pyrazine concentration and larval time in the two sexes.

### Pheromones and female colour: implications for quality signalling

We detected a significant relationship between the amount of pheromone and female colour. Females with redder hindwings produced a higher amount of pheromone. This may indicate that female colouration reflects condition or quality, consistent with previous findings that redder females, on average, develop faster, lay more eggs, and achieve greater weight (Koch et al. 2024). In our data set, we also detected a significantly positive association between redness of female bodies colour and pupa weight (estimate = 169.06 ± 35.6, P = 3.19e-06, Figure S10). There is evidence that producing dark red pigmentation is metabolically costly (Lindstedt et al. 2010), as females subjected to diets with high excretion costs developed paler wings. It seems plausible that only females in good condition, with sufficient resources, can afford to produce the most intense red pigmentation. In this context, pheromones may thus serve as an honest signal to males allowing them to find the highest quality females (i.e. those with the reddest coloration). However, pheromone amount and composition were not directly correlated with female weight, a reliable proxy for reproductive success. Additionally, we found no association between female colour and chemical defence, suggesting that warning colouration does not necessarily serve as an honest signal, i.e. indicating the level of defence (Summers et al. 2015), to predators. Both defence (Burdfield-Steel et al. 2019; Ottocento et al. 2024) and wing colour (Lindstedt et al. 2010), can change plastically in response to diet quality indicating the potential for female colour to act as an honest signal when both traits positively correlate with the amount of a shared resource (Blount et al. 2012). However, this effect may only be detectable if variation in resource availability is sufficiently high, which is unlikely under the controlled and consistent conditions used in this study.

## Conclusion

In addition to the previously known male colour locus, we identified large-effect loci for melanisation, female, and larval warning colour, all mapping to different genomic regions, suggesting that these traits can evolve independently. We found no evidence that male colour is part of a complex phenotype. It is linked to a locus controlling hindwing melanisation, but recombination still occurs. Increased melanisation was associated with reduced activity at high temperatures. We detected some indication of costs related to chemical defence, but no correlation between defence and warning colour. Together, these findings suggest a largely modular genetic basis of warning signal variation in *A. plantaginis*, with different components of the phenotype potentially responding independently to ecological and sexual selection, thereby contributing to the maintenance of colour polymorphism in natural populations.

## Data availability

Sequence data have been deposited at the European Nucleotide Archive under project accession number PRJEB76231. Phenotypic data used for the genetic associations is available from Zenodo (https://doi.org/10.5281/zenodo.20305815). Phenotypic data for the entire laboratory population: https://doi.org/10.5281/zenodo.14018476.

## Author contribution

JM and CDJ conceived the study and provided funding. JM initiated the project and coordinated the maintenance of the lab populations. MNB coordinated the data collection and analysed the images. MNB and JAG collected data and maintained the population. CO collected defence fluids and pheromones and analysed pyrazine concentrations in the defence fluids. CDP collected and analysed pheromones. ES and JM conducted the male activity assay. ON generated the larva photos. SM wrote the image analysis scripts. YFC and MK developed and performed haplotagging, sequencing and sequencing data pre-processing. ELK led the data analysis and wrote the first draft of the manuscript. All authors agreed to the final version of this manuscript.

## Acknowledgments

We thank Kaisa Suisto, Elisa Salmivirta, Jimi Kirvesoja, Liam Murphy and Teemu Tuomaala for their help in maintaining the moth populations during this project. Pheromone extractions were carried out at the Amsterdam pheromone lab.

