## Supporting_Information for "Independent major-effect loci and limited trait associations characterise warning colour variation across sexes and life stages in the wood tiger moth (*Arctia plantaginis*)"

**SUPPORTING INFORMATING**

**Tables**

**Table S1** Sample sizes for testing associations between traits. These may slightly differ from those used for QTL mapping in cases where genotyping failed. For trait explanations see Table 2 and main text.

**Table S2** Results of a quantitative trait loci (QTL) analysis when using genetic data with missing genotypes imputed by STITCH. Each family was analysed separately as a 4-way cross. LG = linkage group; CI = confidence interval. We used grandparents for phasing for families 15, 28, 38, but not for 24 and 25. Positions on linkage groups are therefore not comparable between these two groups of families.

**Table S3** Protein blast results of candidate genes for wing colour and patterns. Those on linkage groups with significant QTLs are highlighted.

**Table S4** Results of testing the effects of male colour genotype and melanism type as well as male colour genotype and melanin genotype on different traits. Linear mixed models with family included as random effect were used. For testing the effects of genotypes, sex was included as fixed effect since we could include females in this test. This was not possible for testing combined effects of colour genotype and melanism since melanism is a male specific trait. Significant results are highlighted. SNPs with the strongest association in the genome wide association studies for male colour and melanism type were used as male colour and melanism genotypes.

**Table S5** Results (Type III ANOVA) for testing the effects of melanisation and male colour on behaviour. Weather (factor: cloudy, half cloudy, sunny/half cloudy, sunny) and temperature were included as fixed and family as random effect. Males were observed for 50 minutes and different activities recorded: time_flipped = minute when the male flipped from its back onto its stomach, A= antennae movement, K = walking, R = fluttering the wings, L = flying, M = other movement. n_active_mins = the number of minutes when any form of activity was observed. sum_active_mins = the sum of all activities observed (i.e. when three different activities were observed within the same minute, this is counted as three minutes). Significant p-values are shown in bold.

**Table S6** Results of testing the effects of pyrazine amount in the defensive fluids on other traits and the effect of pheromone amount and composition. We tested effects of the major defence compounds SBMP (2-sec-butyl-3-methoxypyrazine) and IBMP (2-isobutyl-3-methoxypyrazine), their combined concentration, and the IBMP-SBMP-ratio. For pheromones, we tested the total amount of pheromone and the P6-P3-ratio. Concentrations and ratios of pyrazines and pheromones were log-transformed. We used separate models to estimate the effects of single compounds, the combined concentration and the ratio. Significant effects are highlighted.

### Figures

**Figure S1** Comparison of grouping SNPs into linkage groups (LGs) between a previous data set (Brien et al. 2022) and our new analysis. 0 indicates unassigned SNPs. Assignment to LGs is very consistent. Most SNPs that were assigned to one LG before, were also grouped into one LG in our analysis and there is a clear correspondence between the previous and the new LGs. Inconsistencies concern the longest new LG (LG 1), which had been separated into five LGs (LG 3, 8, 11, 15, 25) before. Another inconsistency is found in the old LG 15, which corresponds partly to our new LG 1, whereas another part (highlighted in red) is grouped together with SNPs of the old LG 12 (LG 7 in our new assignment). Closer inspection showed that all of these SNPs belonged to one scaffold. A quantitative trait analysis (Figure S2) showed that male hindwing colour, which is controlled by one locus on LG 12, showed strong associations with LG 12 as well as with LG 15. This strongly suggest a misassignment of this scaffold, which we modified.

Previous Linkage Groups (columns); New Linkage Group assignment (rows)

| | 0 | 1 | 2 | 3 | 4 | 5 | 6 | 7 | 8 | 9 | 10 | 11 | 12 | 13 | 14 | 15 | 16 | 17 | 18 | 19 | 20 | 21 | 22 | 23 | 24 | 25 | 26 | 27 | 28 | 29 | 30 | 31 |
|---|---|---|---|---|---|---|---|---|---|---|---|---|---|---|---|---|---|---|---|---|---|---|---|---|---|---|---|---|---|---|---|---|
| 0 | 2441 | 3734 | 6500 | 1786 | 10118 | 12786 | 3306 | 6777 | 18965 | 4207 | 5382 | 11276 | 3961 | 5610 | 1157 | 24647 | 19676 | 14788 | 8462 | 25322 | 15446 | 10442 | 1194 | 3143 | 6414 | 3612 | 13781 | 1957 | 1897 | 702 | 4340 | 2346 |
| 1 | 16 | 2 | 2 | 2823 | 9 | 55 | 1 | 6 | 23481 | 4 | 14 | 33158 | 6 | 0 | 1 | 21152 | 5 | 25 | 1 | 6 | 1 | 6 | 0 | 0 | 0 | 9685 | 32 | 3 | 0 | 0 | 0 | 0 |
| 2 | 0 | 0 | 0 | 0 | 0 | 0 | 0 | 0 | 0 | 0 | 0 | 0 | 0 | 0 | 0 | 0 | 0 | 35780 | 0 | 0 | 0 | 0 | 0 | 0 | 0 | 0 | 0 | 0 | 0 | 0 | 0 | 0 |
| 3 | 20 | 0 | 0 | 0 | 0 | 0 | 0 | 0 | 0 | 0 | 0 | 0 | 0 | 0 | 0 | 0 | 0 | 0 | 0 | 31819 | 0 | 0 | 0 | 0 | 0 | 0 | 0 | 0 | 0 | 0 | 0 | 0 |
| 4 | 15 | 0 | 0 | 0 | 0 | 0 | 0 | 0 | 0 | 0 | 0 | 0 | 0 | 0 | 0 | 0 | 0 | 0 | 0 | 0 | 30808 | 0 | 0 | 0 | 0 | 0 | 0 | 0 | 0 | 0 | 0 | 0 |
| 5 | 43 | 0 | 0 | 0 | 0 | 0 | 0 | 0 | 0 | 0 | 0 | 0 | 0 | 0 | 0 | 0 | 0 | 0 | 26296 | 0 | 0 | 0 | 0 | 0 | 0 | 0 | 0 | 0 | 0 | 0 | 0 | 0 |
| 6 | 0 | 0 | 0 | 0 | 0 | 0 | 0 | 0 | 0 | 0 | 0 | 0 | 0 | 0 | 0 | 0 | 0 | 0 | 0 | 0 | 0 | 25383 | 0 | 0 | 0 | 0 | 0 | 0 | 0 | 0 | 0 | 0 |
| 7 | 0 | 0 | 0 | 0 | 0 | 0 | 0 | 0 | 0 | 0 | 0 | 0 | 6425 | 0 | 0 | 16382 | 0 | 0 | 0 | 0 | 0 | 0 | 0 | 0 | 0 | 0 | 0 | 0 | 0 | 0 | 0 | 0 |
| 8 | 0 | 0 | 0 | 0 | 0 | 0 | 0 | 0 | 0 | 0 | 21815 | 0 | 0 | 0 | 0 | 0 | 0 | 0 | 0 | 0 | 0 | 0 | 0 | 0 | 0 | 0 | 0 | 0 | 0 | 0 | 0 | 0 |
| 9 | 0 | 20181 | 0 | 0 | 0 | 0 | 0 | 0 | 0 | 0 | 0 | 0 | 0 | 0 | 0 | 0 | 0 | 0 | 0 | 0 | 0 | 0 | 0 | 0 | 0 | 0 | 0 | 0 | 0 | 0 | 0 | 0 |
| 10 | 1 | 0 | 0 | 0 | 0 | 18914 | 0 | 0 | 0 | 0 | 0 | 0 | 0 | 0 | 0 | 0 | 0 | 0 | 0 | 0 | 0 | 0 | 0 | 0 | 0 | 0 | 0 | 0 | 0 | 0 | 0 | 0 |
| 11 | 0 | 0 | 0 | 0 | 0 | 0 | 0 | 0 | 0 | 0 | 0 | 0 | 0 | 0 | 0 | 0 | 0 | 0 | 0 | 0 | 0 | 0 | 0 | 0 | 17586 | 0 | 0 | 0 | 0 | 0 | 0 | 0 |
| 12 | 11 | 0 | 0 | 0 | 17032 | 0 | 0 | 0 | 0 | 0 | 0 | 0 | 0 | 0 | 0 | 0 | 0 | 0 | 0 | 0 | 0 | 0 | 0 | 0 | 0 | 0 | 0 | 0 | 0 | 0 | 0 | 0 |
| 13 | 0 | 0 | 0 | 0 | 0 | 0 | 0 | 16612 | 0 | 0 | 0 | 0 | 0 | 0 | 0 | 0 | 0 | 0 | 0 | 0 | 0 | 0 | 0 | 0 | 0 | 0 | 0 | 0 | 0 | 0 | 0 | 0 |
| 14 | 0 | 0 | 0 | 0 | 0 | 0 | 0 | 0 | 0 | 0 | 0 | 0 | 0 | 0 | 0 | 0 | 0 | 0 | 0 | 0 | 0 | 0 | 0 | 0 | 0 | 0 | 16196 | 0 | 0 | 0 | 0 | 0 |
| 15 | 1 | 0 | 13041 | 0 | 0 | 0 | 0 | 0 | 0 | 0 | 0 | 0 | 0 | 0 | 0 | 0 | 0 | 0 | 0 | 0 | 0 | 0 | 0 | 0 | 0 | 0 | 0 | 0 | 0 | 0 | 0 | 0 |
| 16 | 0 | 0 | 0 | 0 | 0 | 0 | 0 | 0 | 0 | 0 | 0 | 0 | 0 | 0 | 0 | 0 | 12711 | 0 | 0 | 0 | 0 | 0 | 0 | 0 | 0 | 0 | 0 | 0 | 0 | 0 | 0 | 0 |
| 17 | 0 | 0 | 0 | 0 | 0 | 0 | 0 | 0 | 0 | 12060 | 0 | 0 | 0 | 0 | 0 | 0 | 0 | 0 | 0 | 0 | 0 | 0 | 0 | 0 | 0 | 0 | 0 | 0 | 0 | 0 | 0 | 0 |
| 18 | 0 | 0 | 0 | 0 | 0 | 0 | 0 | 0 | 0 | 0 | 0 | 0 | 0 | 0 | 0 | 0 | 0 | 0 | 0 | 0 | 0 | 0 | 0 | 0 | 0 | 0 | 0 | 0 | 0 | 0 | 6673 | 0 |
| 19 | 1 | 0 | 0 | 0 | 0 | 0 | 0 | 0 | 0 | 0 | 0 | 0 | 0 | 6381 | 0 | 0 | 0 | 0 | 0 | 0 | 0 | 0 | 0 | 0 | 0 | 0 | 0 | 0 | 0 | 0 | 0 | 0 |
| 20 | 5 | 1132 | 0 | 0 | 691 | 0 | 0 | 0 | 0 | 0 | 0 | 0 | 0 | 0 | 0 | 0 | 0 | 0 | 0 | 0 | 0 | 0 | 0 | 0 | 0 | 0 | 0 | 0 | 0 | 0 | 0 | 3339 |
| 21 | 3 | 0 | 0 | 0 | 0 | 0 | 0 | 0 | 0 | 0 | 0 | 0 | 0 | 0 | 0 | 0 | 0 | 0 | 0 | 0 | 0 | 0 | 0 | 0 | 0 | 0 | 0 | 0 | 5105 | 0 | 0 | 0 |
| 22 | 34 | 0 | 0 | 0 | 0 | 0 | 0 | 0 | 0 | 0 | 0 | 0 | 0 | 0 | 0 | 0 | 0 | 0 | 0 | 2 | 0 | 0 | 0 | 0 | 0 | 0 | 0 | 5002 | 0 | 0 | 0 | 0 |
| 23 | 0 | 0 | 0 | 0 | 0 | 0 | 0 | 0 | 0 | 0 | 0 | 0 | 0 | 0 | 0 | 0 | 0 | 0 | 0 | 0 | 0 | 0 | 0 | 4927 | 0 | 0 | 0 | 0 | 0 | 0 | 0 | 0 |
| 24 | 5 | 650 | 0 | 0 | 0 | 0 | 0 | 0 | 0 | 0 | 0 | 0 | 0 | 0 | 3067 | 0 | 0 | 0 | 0 | 0 | 0 | 0 | 0 | 0 | 0 | 0 | 0 | 0 | 0 | 0 | 0 | 0 |
| 25 | 27 | 0 | 0 | 0 | 0 | 0 | 3630 | 0 | 0 | 0 | 0 | 0 | 0 | 0 | 0 | 0 | 0 | 0 | 0 | 0 | 0 | 0 | 0 | 0 | 0 | 0 | 0 | 0 | 0 | 0 | 0 | 0 |
| 26 | 0 | 0 | 0 | 0 | 0 | 0 | 0 | 0 | 0 | 0 | 0 | 0 | 0 | 0 | 0 | 0 | 0 | 0 | 0 | 0 | 0 | 0 | 2179 | 0 | 0 | 0 | 0 | 0 | 0 | 0 | 0 | 0 |
| 27 | 49 | 0 | 0 | 0 | 0 | 0 | 0 | 0 | 0 | 0 | 0 | 0 | 0 | 0 | 0 | 0 | 0 | 0 | 0 | 0 | 0 | 0 | 0 | 0 | 0 | 0 | 0 | 0 | 0 | 1354 | 0 | 0 |

**Figure S2** Preliminary results of a quantitative trait loci analysis (A) and a genome wide association analysis (B) for male hindwing colour. This trait is controlled by one locus on linkage group (LG)12 (Brien et al. 2022). However, due to an error in the previous assignments of SNPs to linkage group, an additional association is found on LG 15. We corrected this in our new linkage map.

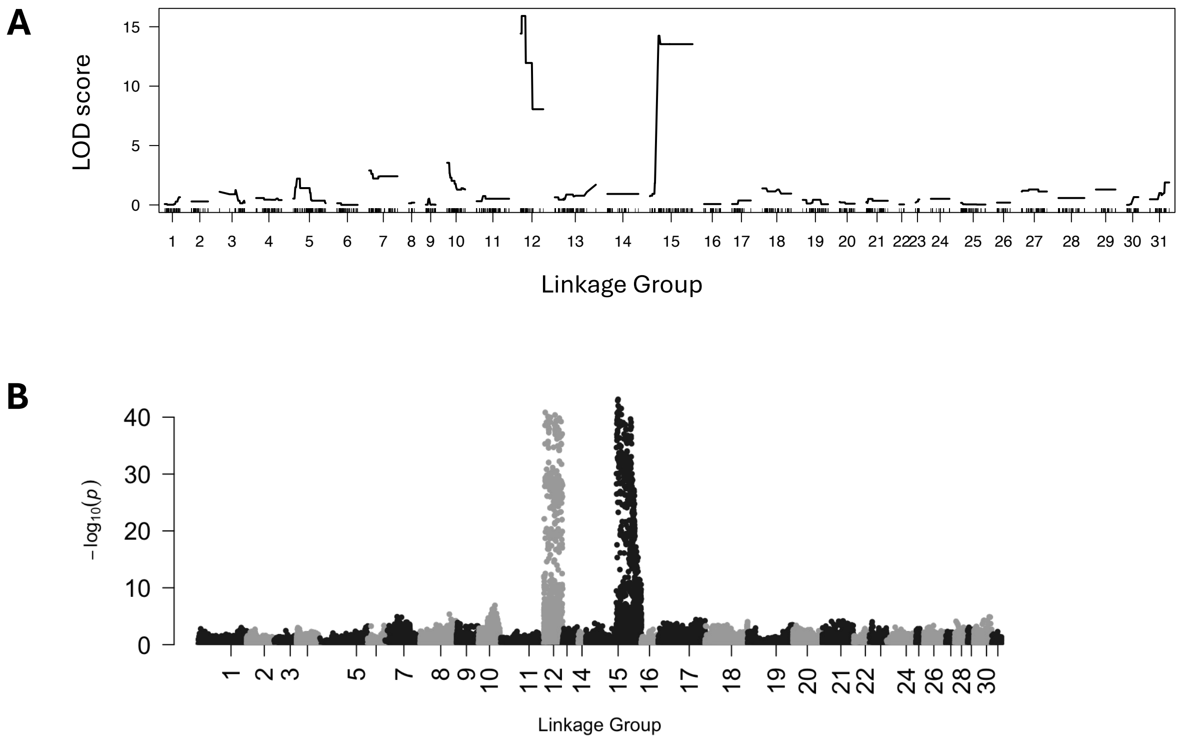


**Figure S3** Average sequencing depth for each LG in females, N = 321 (A) and males, N = 309 (B). The ratio of average female depth: male depth for each scaffold is shown in C. LG 0 indicates scaffolds that could not be assigned to any LG.

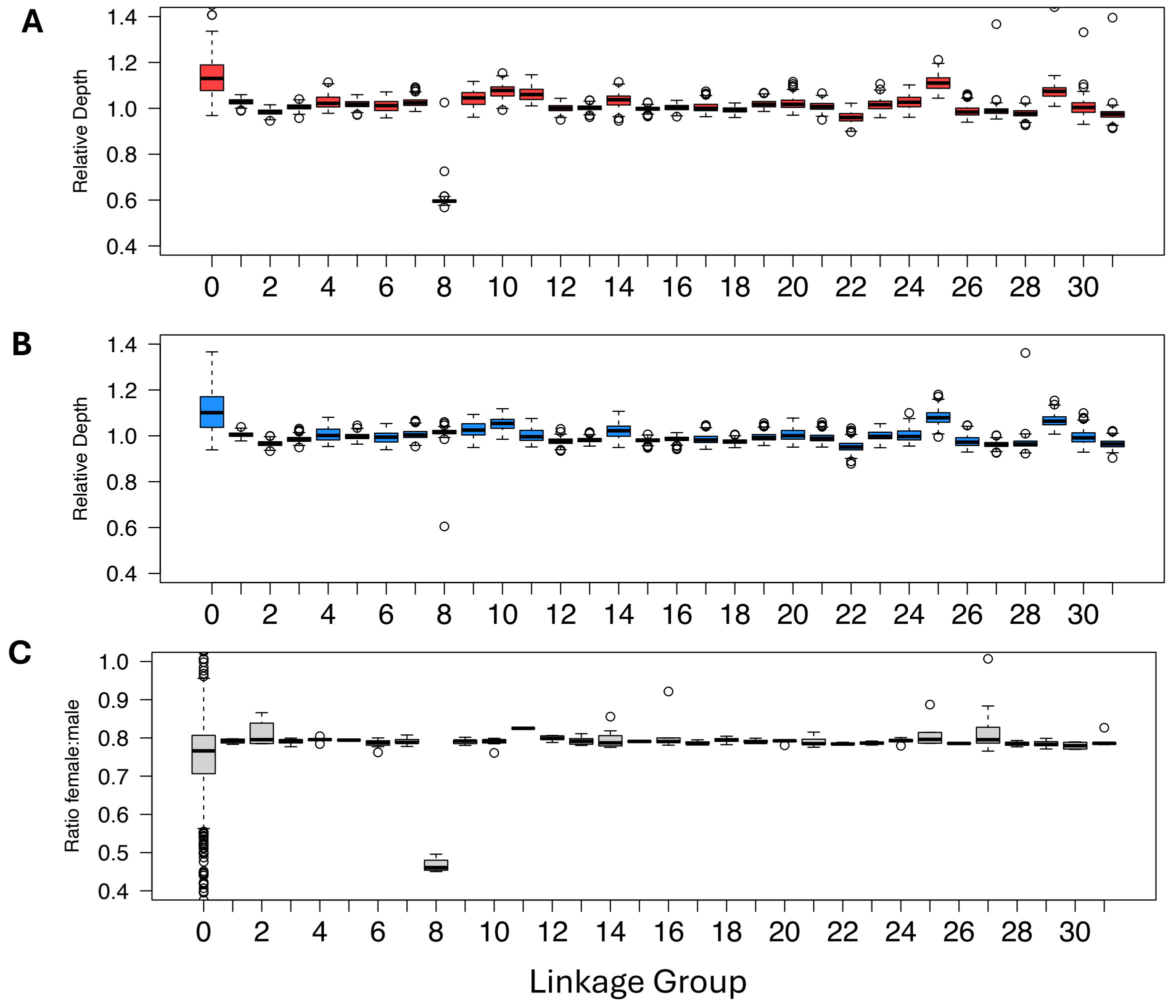


**Figure S4** Genome wide association (GWAS) with sex as binary trait. The highly significant region at the end of LG11 showed a sequencing depth that was much higher in females than in males, and within females much higher than the average suggesting that this part of a misassembled W-chromosome. LG 8 was identified as Z-chromosome based on sequencing depth (see Figure S3). Red line indicates -log10(5e-8).

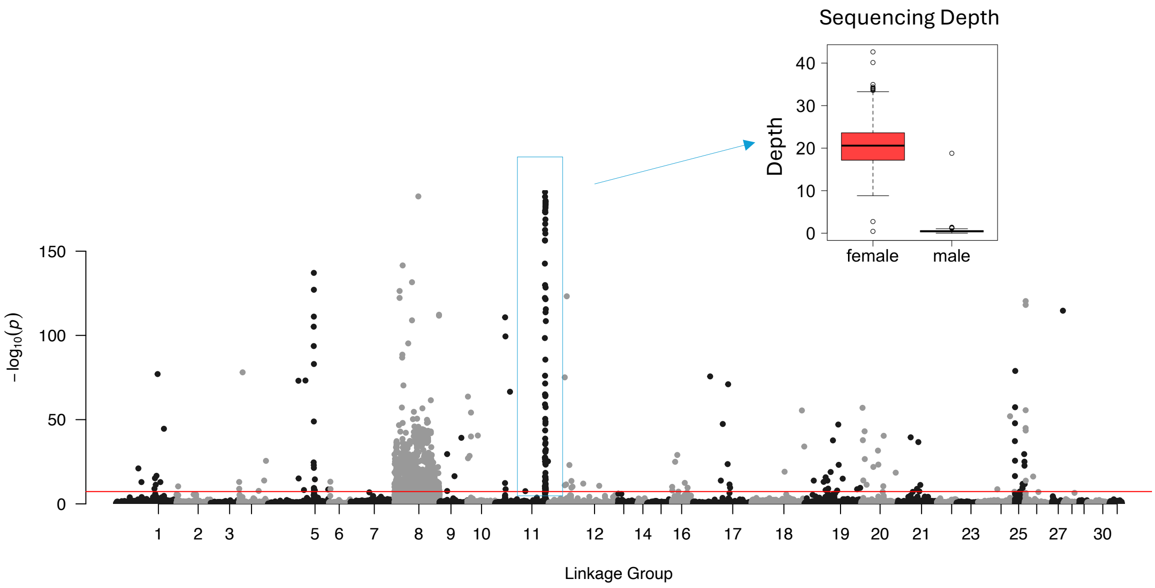


**Figure S5** Larval developmental times in the laboratory population over time in white and yellow morphs (A). Proportion of the plus-melanism type within each colour morph in the laboratory population over time (B). Comparison of larval times between plus and minus melanisation morphs (C). Larval times of overwintering individuals were excluded.

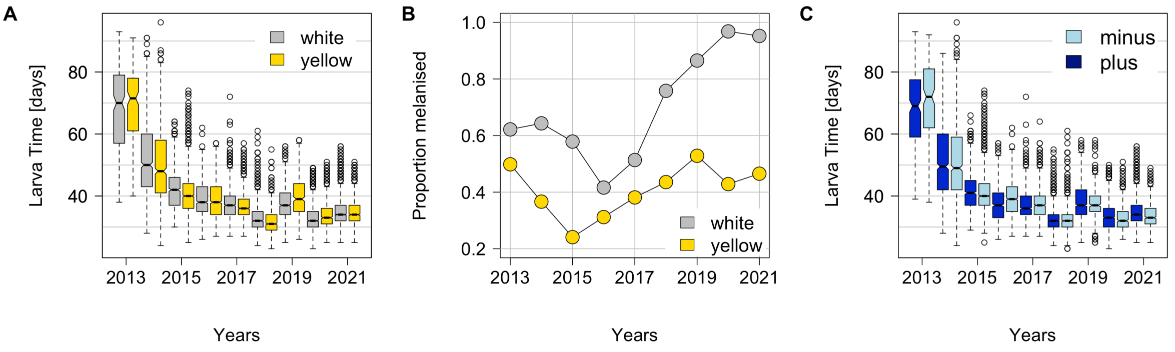


**Figure S6** GWAS for melanism type (plus or minus as binary trait) using all males (A) and yellow males only (C). B (all) and D (yellows only) show associations for SNPs on LG 12 with the positions of candidate genes *cortex, domeless, washout* (Table S3) indicated by the red line.

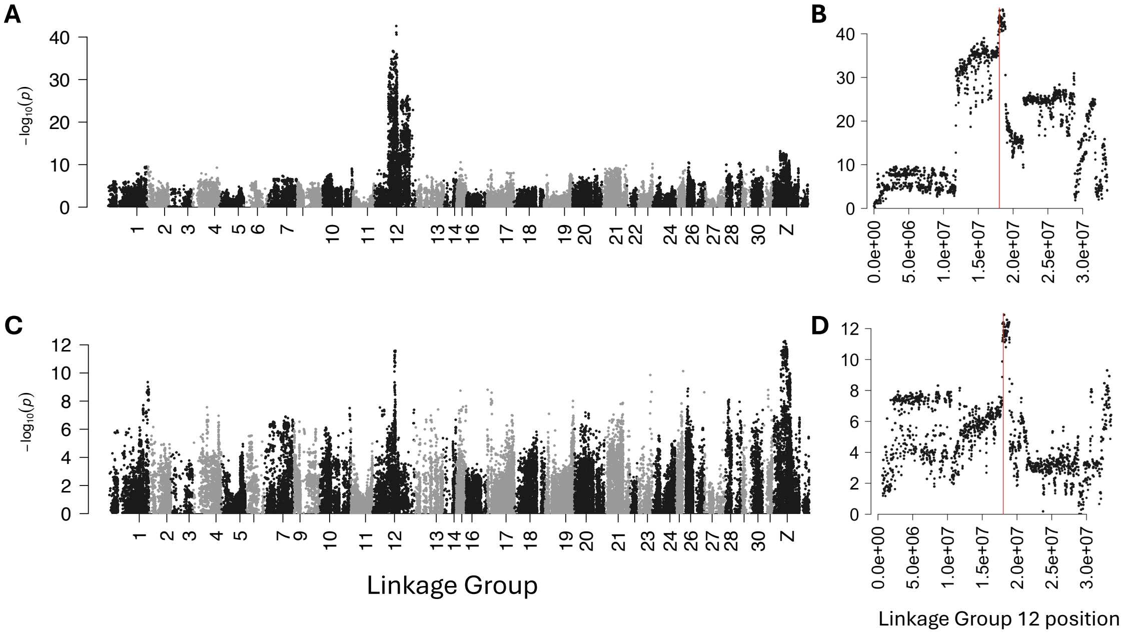


**Figure S7** Effect of weather on activity behaviour in males (A). Males were observed for 50 minutes. Shown are the number of minutes when any form of activity was observed. B: Changes of activity with increasing temperature. Shown are observations and model predictions with confidence intervals. The model (linear mixed model) included weather, temperature, and proportion of melanised hindwings as fixed effects and family as a random effect.

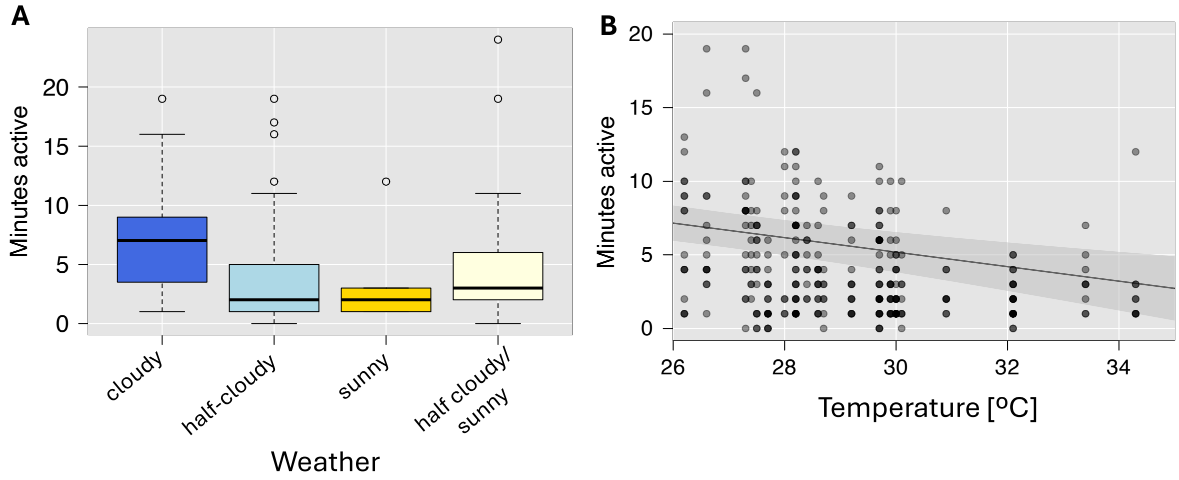


**Figure S8** Effects of increasing temperature on the activity behaviour of males with weakly (light blue) and strongly (darker blue) melanised hindwings. The inset shows the distribution of the proportion of melanised hindwings. Those that were classified as weakly/strongly melanised are highlighted.

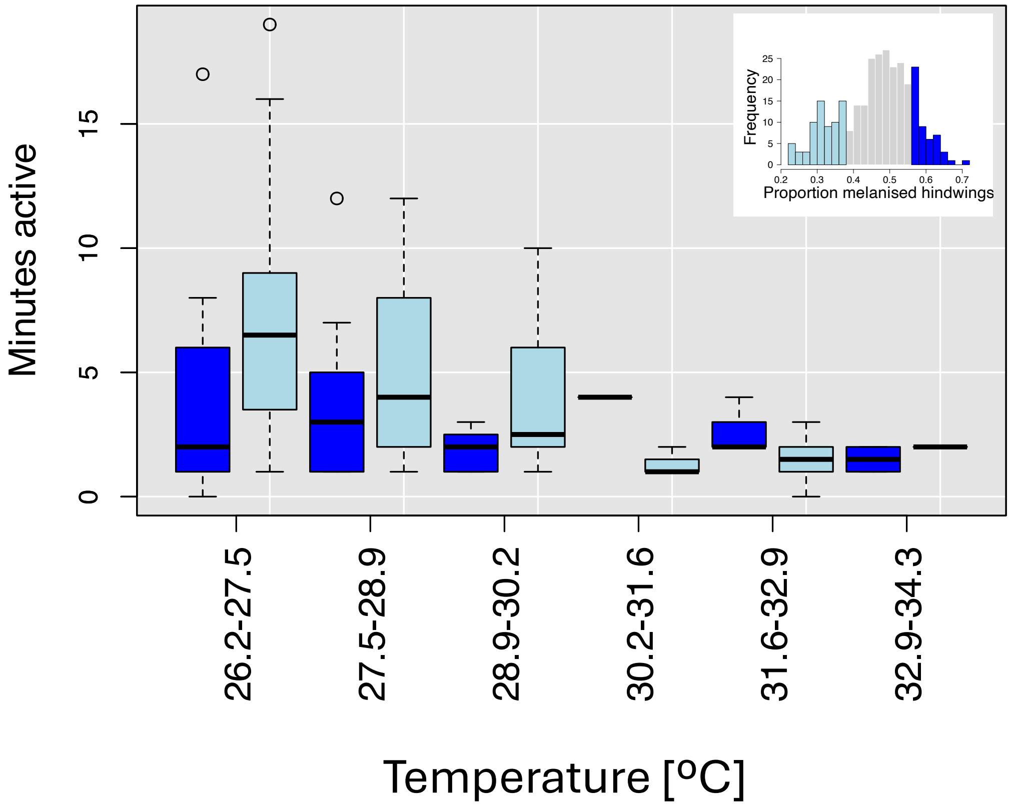


**Figure S9** GWAS for larval signal size (see main Figure 3I) on linkage group 21. The red line indicates the position of the gene *optix*.

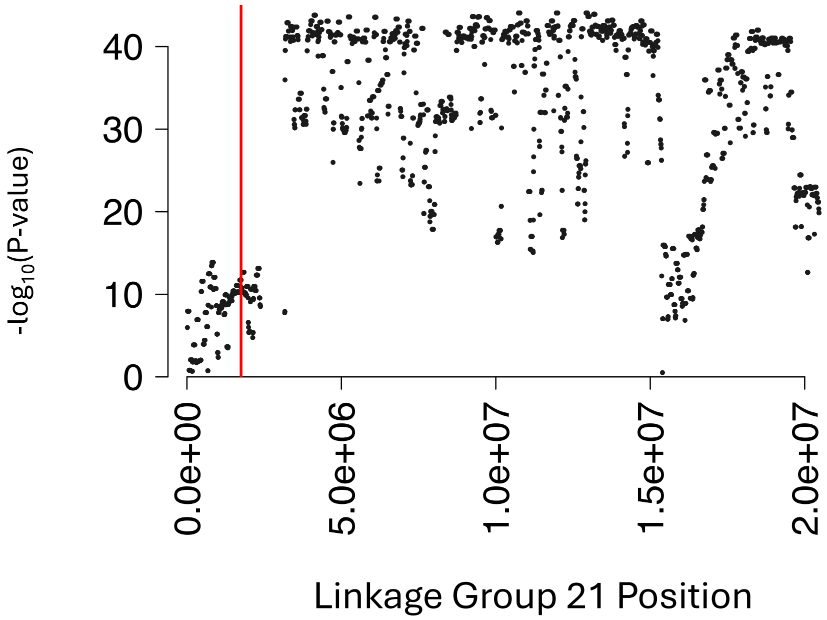

**Figure S10** Relationship between female body colour (‘rg_body’) quantified from images and pupa weight. Lower rg_body values indicate a yellow colour, higher values a red colour. Shown are also predictions and confidence intervals from linear mixed models including family as a random effect.

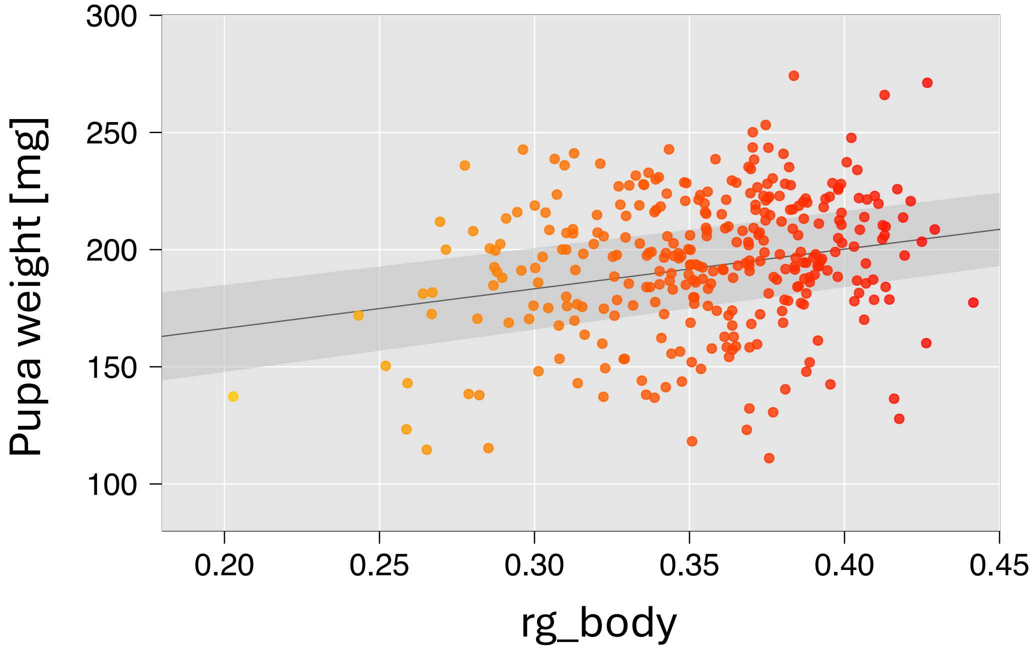


**Appendix**

*Image Analysis*

Photographs were taken of late-stage larvae from above, and adult wings and bodies separately. All photographs were taken under standard lighting conditions with a Samsung NX1000 digital camera converted to full-spectrum with no quartz filter to enable ultraviolet sensitivity. A UV and infrared blocking filter was used for the human-visible photos, which transmits wavelengths between 400 and 680 nm (Baader UV/IR Cut Filter). A UV pass filter (Baader U filter), which transmits wavelengths between 320 and 380 nm, was used for UV images. All images included a grey-scale reflectance standard (Avian Technologies, Micro FSS08). Photos were standardised using the reflectance standard, and scaled using a ruler in the image with custom MATLAB scripts.

For the larvae photographs, an outline was drawn around the body and around the orange patch in MATLAB. The number of pixels in each shape was used to calculate the proportion of orange regions of the larva compared to the whole body. RGB values were extracted from the orange patch and redness calculated.

For the male forewings and hindwings, a Segment Anything Model was used to extract the proportion of melanised (black) regions of the wings compared to the non-melanised (white or yellow) regions (custom MATLAB script). We used the positions of the non-melanised regions to extract RGB values from the same positions in the UV photos, and used the L value as a measure of UV brightness. UV brightness was only measured from male hindwings, since forewings reflect little UV.

Redness of both female hindwings and the red patches on the adult bodies were measured using the Color Threshold function in ImageJ (v1.53), and RGB values extracted from these regions with the Color Histogram plugin.

*Chemical defence fluid*

After eclosion, thoracic fluid was extracted by squeezing the moths' thorax with tweezers and collected in a 10ul glass capillary. The volume was measured with a calliper. Samples were transferred to glass vials and stored at −20°C until analysis. We measured concentration of SBMP (2-sec-butyl-3-methoxypyrazine) and IBMP (2-isobutyl-3-methoxypyrazine), the main defence compounds in the defensive neck fluids released by adults following the methods of Cai et al. (2007) as described in Burdfield-Steel et al. (2018). Prior to GC_MS analysis, samples were thawed and mixed with a 200-µl NaCl solution (3%). Pyrazines were extracted from the headspace of fluid samples using SPME (solid phase microextraction) fibres for 30 min at 37°C. GC/MSD and analyses were run on an Agilent 7890A series GC system equipped with a Zebron ZB-5MSi, GC Cap. column (length 30 m,0.25 mm I.D. with a film thickness of 0.25 µm) connected to a mass spectrometer Agilent 7000 MS. The fibres were manually loaded into the injector using a spitless injection mode, and the inlet temperature was set to 260°C. Helium was used as a carrier gas at a constant flow rate of 0.8 mL/min. The oven temperature was programmed as follows: 3 min at 60°C then ramped to 170°C at a rate of 7°C/min and from 170 to 260°C at a rate of 20°C/min and kept at that temperature for an additional 5 min. SBMP and IBMP were detected using selected ion monitoring of ions 124, 138 and 151. Once the analysis was completed, the chromatograms and mass spectra were evaluated using Agilent Chemstation (v. G1701CA) software and the Wiley8th edition mass spectral database and the methoxypyrazines were identified using the ratio of these detected ions from the NIST web-book page (Stein), as well as by comparison with standards of SBMP and IBMP. The amount in ng of the two methoxypyrazines in the sample was calculated by comparison with known amounts of the standards run in the same manner as the fluid samples. The amounts of IBMP, SBMP, their combined amounts and their ratio were log-transformed before analysis.

*Female sex pheromones*

Female sex pheromones were collected when females were 1-2 days old. The pheromone gland, located at the tip of the abdomen, was excised with surgical scissors and soaked for 45 minutes in 50ul of hexane containing 200ng of n-pentadecane standard used as a known quantity to later quantify the pheromone compounds. The pheromone solution was stored at -20°C until analysis at the University of Amsterdam. Before analysis, the pheromone solution was resuspended in 5ul of hexane for about 30 min. Next, 2ul of the extract, topped with 1ul of octane were injected in the gas chromatography (GC). We measured the amount of two compounds, P3 and P6, by comparison to reference compounds and quantified by integrating the area under the peak in the chromatogram using Agilent ChemStation software. Absolute amounts (in ng) of each compound were calculated relative to the 200 ng of internal standard. Finally, the ratio of the second major compound (P6) was calculated over the amount of the major compound (P3). Measurements of the total pheromone amount as well as P6.P3 ratio were log-transformed.

*Linkage Group assignment*

We used Lep-MAP3 (Rastas 2017) for linkage map construction. After calling parental genotypes with 'ParentCall2' and filtering markers based on segregation distortion, we separated markers into linkage groups (LG) using 'SeparateChromosomes2' with sizeLimit=100, femaleTheta=0 to account for lack of recombination in females, and a range of different values for the lodLimit parameter (5-25) and different data sets as input: the filtered VCF as well as the same filtered VCF with missing genotypes imputed with STITCH using k=8, k=10, and ngen=30 (other parameters as described in the main manuscript) and a VCF with STITCH imputed genotypes with INFO.SCORE > 0.5 (see main manuscript). We obtained the best grouping into LGS, i.e those that is closest to the number of chromosomes (Yen et al. 2020), using the STITCH imputed VCF and lodLimit=12 femaleTheta=0 distortionLod=1 informativeMask=2 sizeLimit=100 for 'SeparateChromosomes2'. We compared this linkage group assignment to a previous one, which was based on samples used in Brien et al. (2023). Since the set of SNPs differed between both data sets, we used the assignments of scaffolds to LGs in the previous data set for comparisons. In most cases, all SNPs from the same scaffold had been assigned to the same LG in the previous analysis and the proportion of inconsistent SNPs, i. e. SNPs from the same scaffold assigned to different LGs, was very small (< 5%). We then compared to which LGs SNPs of each scaffold were assigned in our new analysis and compared this with the previous scaffold-LG assignment. We found that SNPs from the same scaffold were generally assigned to the same LG and in most cases, the grouping of scaffolds into LGs was consistent with the previous one apart from our longest LG 1, which corresponded to different previous LGs. One exception was a group of SNPs on a scaffold assigned to the previous LG 15 which grouped with SNPs assigned to the previous LG 12 in our assignment (see Figure S1). We first used the previously detected LGs to be consistent and to achieve the correct number of LGs, i.e. according to the number of chromosomes. In a first QTL analysis with male colour as a trait, which we used as a positive control to assess the quality of our linkage maps and the phasing of the data, we detected two QTL peaks (on LG 12 and LG 15) for this Mendelian trait (Figure S2). Together with the incongruency in LG assignment this strongly suggests an assignment error, i.e. part of LG 15 belonging to LG12. We identified that all SNPs associated with male colour on LG 15 belonged to the scaffold "WW_tarseq_540_arrow" (positions > 12,261,029). A GWAS analysis of male colour confirmed that SNPs on that scaffold were closely linked to the colour locus (Figure S2), which is known to be on LG

12 (Brien et al.). We modified the LG assignment accordingly and assigned parts of the scaffold (WW_tarseq_540_arrow: positions > 12,261,029) to LG 12.

*Sex Chromosome Identification*

To identify the Z chromosome among the LGs, we compared read depths of all LGs separately in females and males. The LG corresponding to the Z-chromosome should show only half the read depth in females compared to other LGs but no differences in males. We used samtools (Li and Durbin 2009) bedcov to obtain the read depth for all individuals for each LG. Comparing males and females we found obvious differences for LG 8 (Figure S3). Additionally, we tested for an association between sex and genomic regions in a genome wide association analysis (GWAS, details below). SNPs on LG 8 showed an elevated level of association (Figure S4). Furthermore, there was a narrow region at the end of LG 11 with SNPs showing an extremely high association (log P-value >150, see Figure S4) and the respective genomic region revealed an increase read depth in females only suggesting that this part of the genome belonged to the W-chromosome and was miss-assembled. We subsequently removed the corresponding markers from QTL-analysis (scaffold WW_467_tarseq, positions 14,000,000-16,800,000). We also used the ZLimit=2 option in 'ParentCall2' in LepMap to call sex chromosome markers. Markers identified as sex-linked by LepMap were those assigned to LG 8 and LG 11 (within the GWAS-sex peak). 99.8 % of the identified markers belonged to LG 8, 0.2 % to LG 11, scaffold WW_467_tarseq.

*QTL analysis using STITCH generated VCF*

We generated linkage maps and conducted a quantitative trait loci (QTL) mapping analysis using the VCF generated by STITCH with missing genotypes imputed. We kept only the most reliable genotypes (INFO SCORE > 0.5), resulting in 9,062,667 variants. We reduced this data set and kept only every 10$^{th}$ variant. Computational time for generating linkage map scales exponentially with the number of markers. Additionally, there would be no advantage in keeping all SNPs since the resolution in QTL mapping is limited by the number of recombination events and not by the number of genetic markers, especially when the number of samples is small as was the case in our families. Steps for generating the linkage map were the same as described for the filtered VCF without imputed genotypes with the exception of using the option 'phasedData=1' when ordering the markers of each LG using 'OrderMarker2' since STITCH outputs a phased VCF file. After filtering for segregation distortion 737,860 SNPs were left, which were then separated into 31 LGs using the previous Scaffold-LG assignment as described above. Markers were ordered with and without using grandparents for phasing. The QTL mapping was done in the same way as described in the main manuscript. We used the grandparental phased data sets for families 15 and 38, and the data phased without using grandparents for families 24, 25, and 28. The resulting linkage maps consisted of 31 LGs ranging in size from 145.3-316.6 cM with a total length of 6218.67 cM and 7693 unique positions (grandparental phased) and 141.0-294.2 cM with a total length of 6101.40 cM and 7523 unique positions without grandparental phasing.

Results were consistent with those obtained from the VCF without imputed genotypes: There were significant QTL peaks for different measures of female coloration on LG 27 (female colour, rg_wing, rg_body in family 15, rg_body in family 25), for melanism on LG 12 (in family 15 and 38). We also detected a QTL for larva signal size on LG 21 in family 24 and 25 that was only found family 24 in the other analysis (see Table S2).
